# SGC-CAMKK2-1: A chemical probe for CAMKK2

**DOI:** 10.1101/2022.10.18.512752

**Authors:** Carrow Wells, Yi Liang, Thomas L. Pulliam, Chenchu Lin, Dominik Awad, Benjamin Eduful, Sean O’Byrne, Mohammad Anwar Hossain, Carolina Moura Costa Catta-Preta, Priscila Zonzini Ramos, Opher Gileadi, Carina Gileadi, Rafael M. Couñago, Brittany Stork, Christopher G Langendorf, Kevin Nay, Jonathan Oakhill, Debarati Mukherjee, Luigi Racioppi, Anthony Means, Brian York, Donald P. McDonnell, John W. Scott, Daniel E. Frigo, David H. Drewry

## Abstract

The serine/threonine protein kinase calcium/calmodulin-dependent protein kinase kinase 2 (CAMKK2) plays critical roles in a range of biological processes. Despite its importance, only a handful of inhibitors of CAMKK2 have been disclosed. Having a selective small molecule tool to interrogate this kinase will help demonstrate that CAMKK2 inhibition can be therapeutically beneficial. Herein, we disclose SGC-CAMKK2-1, a selective chemical probe that targets CAMKK2.

## 1. Introduction

Kinases have been successfully targeted by medicinal chemistry programs to develop therapies for diseases, mostly directed towards oncology indications. Although there are over 70 approved small molecule inhibitors targeting kinases (www.brimr.org/PKI/PKIs.htm), many of the >500 protein kinases have garnered little attention and have no published selective or potent inhibitors. CAMKK2 (Calcium/Calmodulin-Dependent Protein Kinase Kinase 2) is a kinase that is important in physiological processes such as energy balance[1] and many pathological processes (Table 1) as well. For example, CAMKK2 activation has been linked to several types of cancers, metabolic diseases, and regulation of the immune response (Table 1). Despite having multiple links to disease, and thus a potential therapeutic target, there is only one tool molecule that is commercially available and in widespread use - STO-609 (**Figure 1**).

**Table 1.**
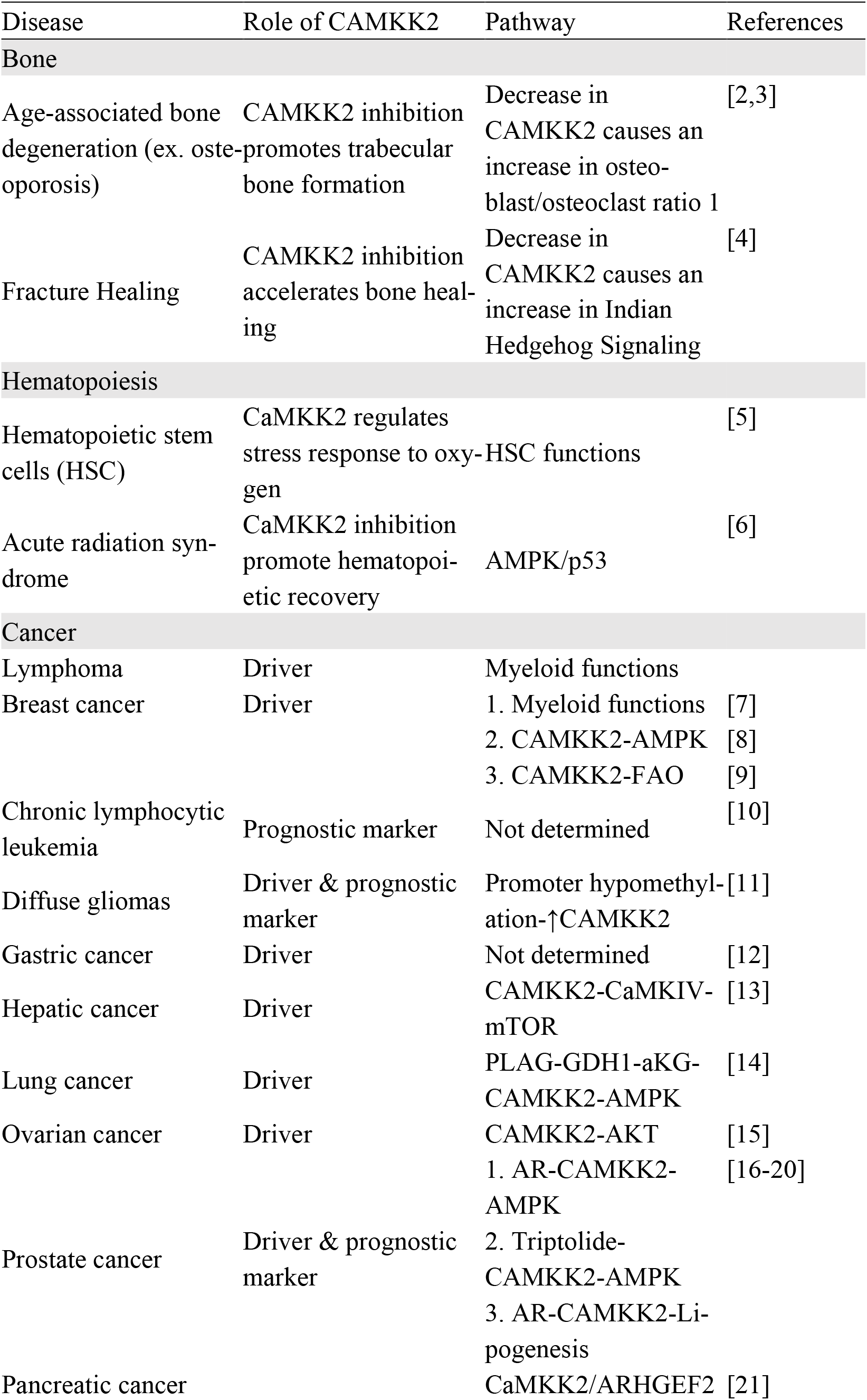
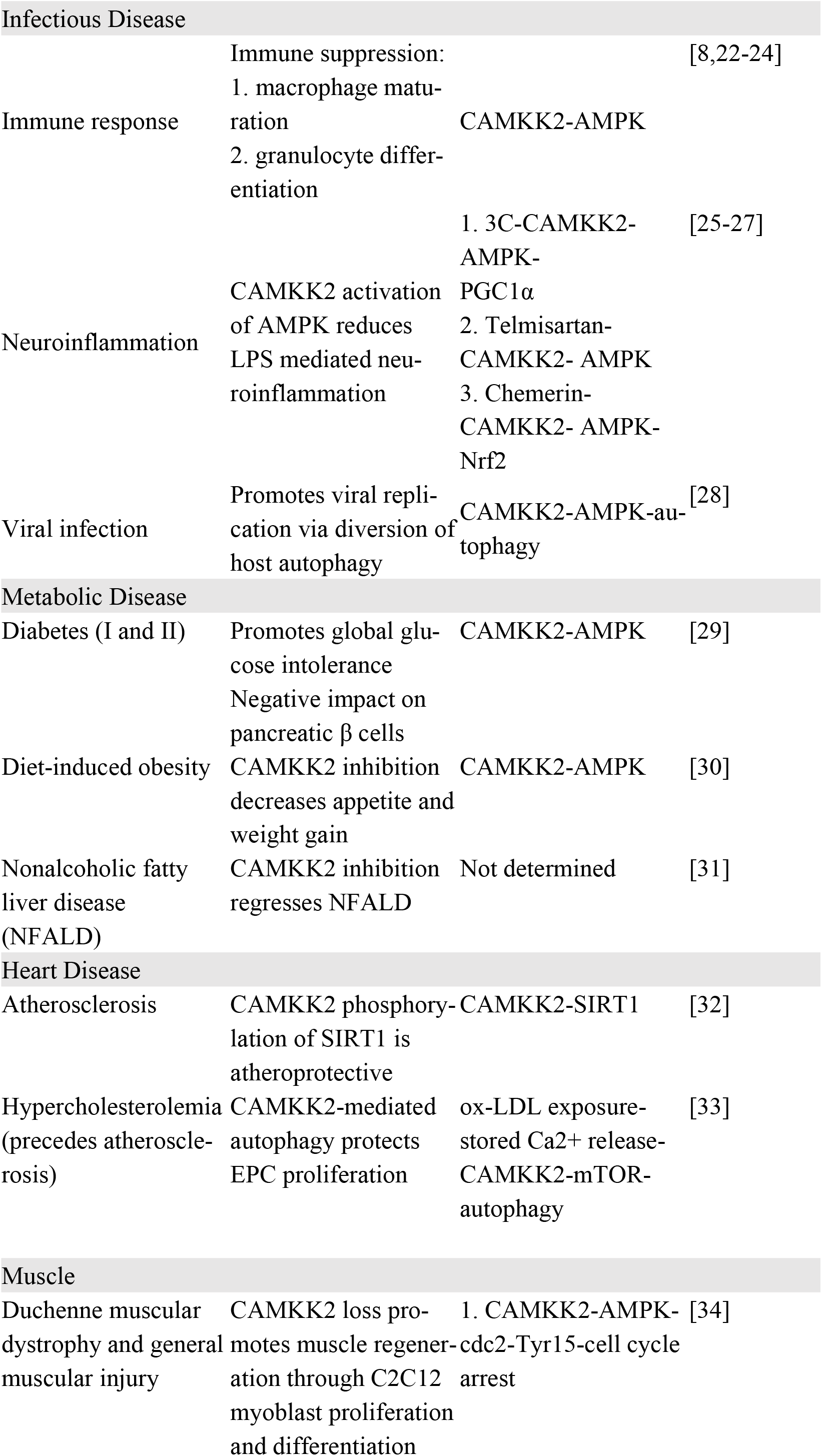
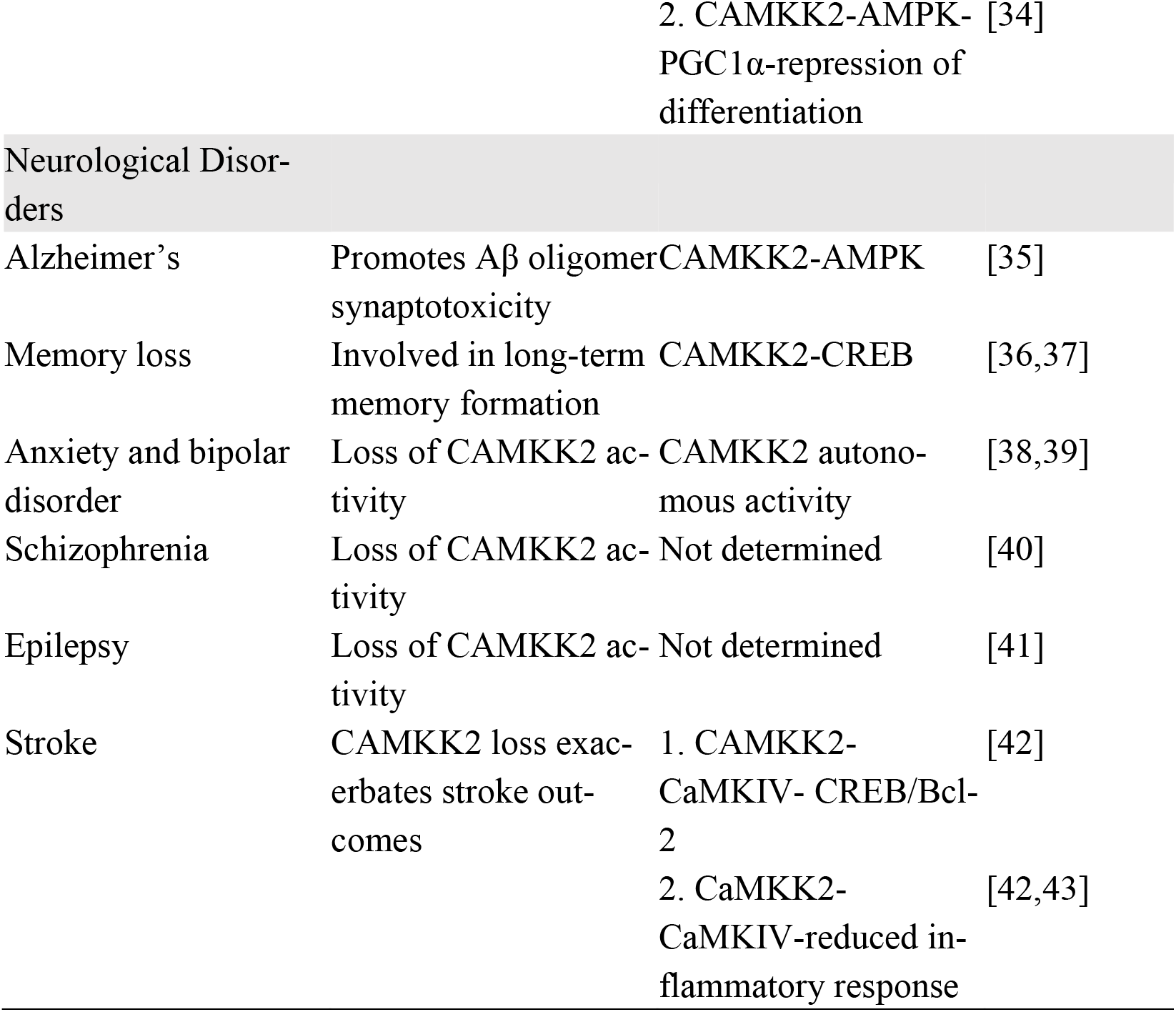
CAMKK2 disease links.

**Figure 1.**
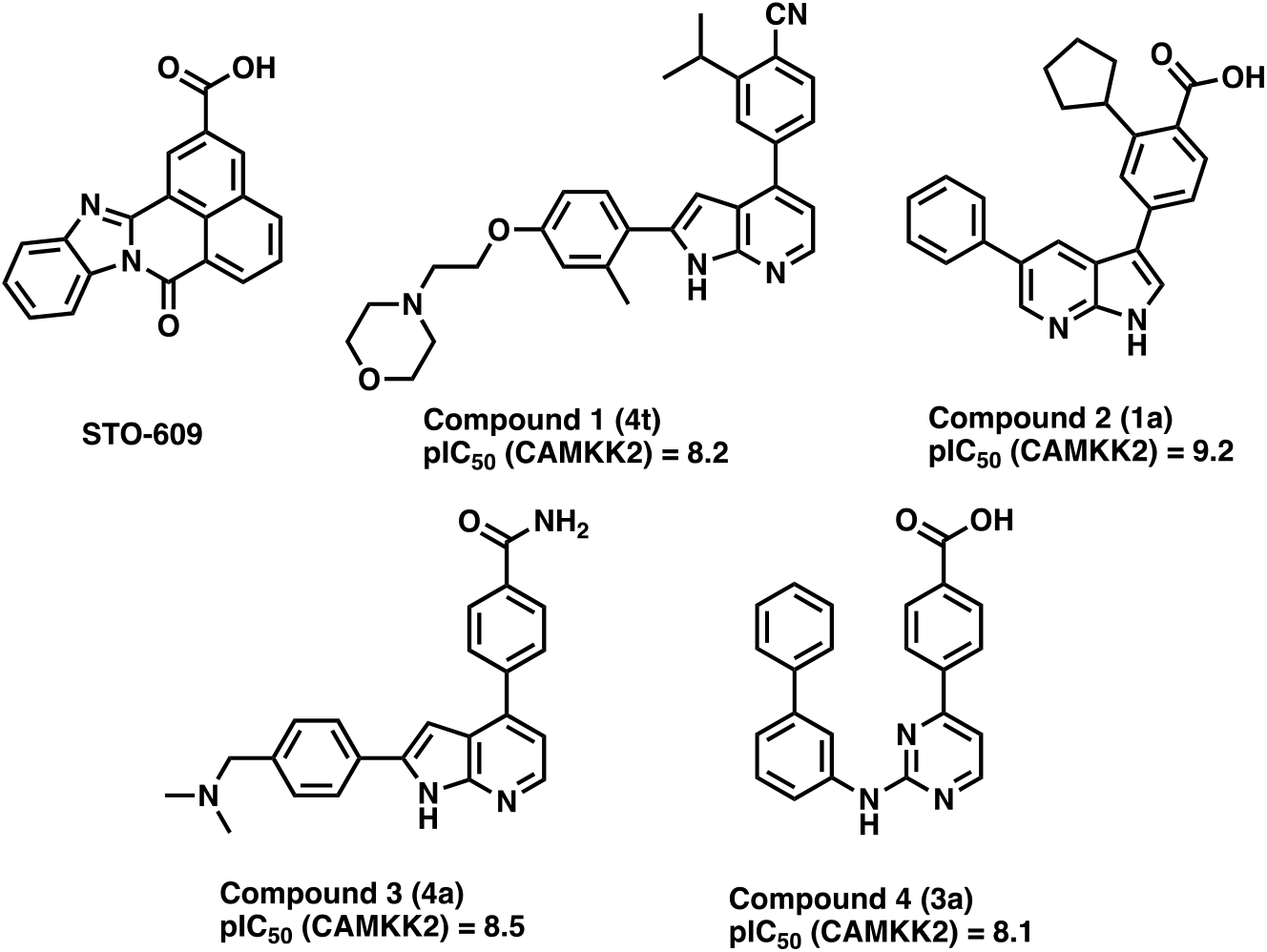
STO-609 and some reported (Price et al. numbering noted in parenthesis [47]) CAMKK2 inhibitors.

The compound STO-609 has demonstrated CAMKK2 inhibition in both cellular and in vivo settings [7,31,44]. However, it has also been shown to inhibit additional kinases such as CK2, AMPK, MNK1, PIM2, PIM3, DYRK2, DYRK3, and ERK8, which at high treatment concentrations could lead to confounding results. Recently, larger screening efforts have been undertaken to identify small molecules defined as inhibitors for other kinases that can inhibit CAMKK2 [45]. These studies were performed using a thermal shift (differential scanning fluorimetry, DSF) and enzyme inhibition assays. From these efforts, 20 compounds stabilized the protein more than 10°C and 10 compounds inhibited CAMKK2 in the enzyme assay at concentrations <100 nM. These compounds, while potent, inhibit additional kinases making any phenotype observed using these compounds difficult to attribute directly to CAMKK2 inhibition. To address this limitation, we sought to develop a potent and selective CAMKK2 probe molecule that could be utilized to investigate the biological implications of CAMKK2 inhibition more thoroughly. A probe molecule is a potent and selective small-molecule inhibitor that has a well-characterized mechanism of action allowing the user to attribute a biological response directly to a target [46]. In addition to the probe molecule, a structurally related but inactive compound should ideally be available to be profiled alongside the probe to give additional confidence in the biological activity of the probe.

A few other promising CAMKK2 inhibitors have been disclosed in the literature, such as a recent publication where scientists from GSK describe three related CAMKK2 inhibitor scaffolds that were optimized to afford a blood-brain penetrant inhibitor, compound **1 (4t) (Figure 1**) [47]. These molecules, despite their single digit nanomolar CAMKK2 potency, were shown to have activity beyond CAMKK2 in the small panel they were evaluated against, making them good starting points for further optimization, but suboptimal for directly interrogating CAMKK2’s roles in biology. To obtain a molecule whose biological results could be more directly attributed to CAMKK2 inhibition, we started a medicinal chemistry campaign to identify a probe using the very potent compound **2** (GSK650394) as a chemical starting point.

## 2. Materials and Methods

### 2.1 Synthesis

#### General chemistry information

All reagents and solvents, unless specifically stated, were used as obtained from their commercial sources without further purification. Solvents were degassed with nitrogen for cross-coupling reactions. Air and moisture sensitive reactions were performed under an inert atmosphere using nitrogen in a previously oven-dried reaction flask, and addition of reagents were done using a syringe. All microwave (μW) reactions were carried out in a Biotage Initiator EXP US 400W microwave synthesizer. Thin layer chromatography (TLC) analyses were performed using 200 μm pre-coated sorbtech fluorescent TLC plates and spots were visualized using UV light. High resolution mass spectrometry samples were analyzed with a ThermoFisher Q Exactive HF-X (ThermoFisher, Bremen, Germany) mass spectrometer coupled with a Waters Acquity H-class liquid chromatograph system. All HRMS were obtained *via* electrospray ionization (ESI). Column chromatography was undertaken with a Biotage Isolera One or Prime instrument. Nuclear magnetic resonance (NMR) spectrometry was run on a varian Inova 400 MHz or Bruker Avance III 700 MHz spectrometer equipped with a TCI H-C/N-D 5 mm cryoprobe and data was processed using the MestReNova processor. Chemical shifts are reported in ppm with residual solvent peaks referenced as internal standard.

### 2.2 Eurofins DiscoverX Broad Kinome Profiling (KINOME*_scan_*™)

Compounds were screened at Eurofins DiscoverX (Freemont, CA) at a single concentration of 1 μM using binding assays as described previously [48,49]. Briefly, extracts containing DNA-tagged kinases are incubated with immobilized kinase inhibitors and the test compound (at 1 μM). Test compounds that displace the immobilized broad-spectrum inhibitors are detected by quantitative PCR. The results are expressed as the percentage of kinase bound to the bead compared to DMSO control (%control). Compounds with high affinity have %control of <10. A measure of selectivity is the S_10_ (1 μM) which provides a measure of kinases that demonstrate PoC<10 at a particular concentration, in this case 1 μM.

S10 Calculation

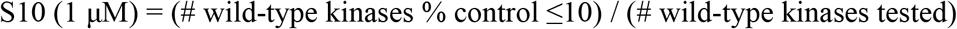

### 2.3 CAMKK2 Enzyme Assay

CAMKK2 activity was determined by measuring the transfer of radiolabeled phosphate from [γ-32P]-ATP to a synthetic peptide substrate (CaMKKtide) as previously described [50]. Briefly, purified recombinant CAMKK2 (100 pM) was incubated in assay buffer (50 mM HEPES [pH 7.4], 1 mM DTT, 0.02% [v/v] Brij-35) containing 200 μM CaMKKtide (Genscript), 100 μM CaCl2, 1 μM CaM (Sigma-Aldrich, Castle Hill, NSW, Australia) 200 μM [γ-32P]-ATP (Perkin Elmer, Boston, MA, USA), 5 mM MgCl2 (Sigma-Aldrich, Castle Hill, NSW, Australia) and various concentrations of inhibitors (0–1 μM) in a standard 30 μl assay for 10 min at 30 °C. Reactions were terminated by spotting 15 μl onto P81 phosphocellulose paper (GE Lifesciences, Paramatta, NSW, Australia) and washing extensively in 1% phosphoric acid (Sigma-Aldrich, Castle Hill, NSW, Australia). Radioactivity was quantified by liquid scintillation counting.

### 2.4 NanoBRET Cellular Target Engagement

CAMKK2 NanoBRET assay: To quantify the cellular activity of these inhibitors we developed a CAMKK2 NanoBRET target engagement assay [51,52]. Briefly this assay utilizes a nanoluciferase (NL) fused to the kinase domain of CAMKK2. This NL kinase fusion is then transiently transfected into HEK293 cells and after 24 hours, tracer is added to the cells. When the tracer and the NL-CAMKK2 fusion come into proximity they create a BRET signal that can be competed in a dose-dependent manner by the addition of cell-penetrant CAMKK2 inhibitors.

### 2.5 In silico docking of SGC-CAMKK2-1

SGC-CAMKK2-1 was docked onto the available crystal structure of CAMKK2 bound to a similar furo-pyridine compound (13g from PMID: 34264658/ PDB ID 5UY6)[52]. Co-crystal structure coordinates were downloaded from the Protein DataBank and prepared for docking using the protein preparation workflow in Maestro (Schrödinger. LLC, New York, USA; version 13.0.137) using default settings [53]. Briefly, missing side chains for residues Glu277, Glu361, Arg363 and Glu439 were filled in (no atoms in these residues were < 10 Å from the ligand); missing residues (214-221) were not filled in and protein termini were not capped; potential hydrogen bond assignments were optimized using PROPKA [54]; energy minimization was performed (heavy atoms were restrained with a harmonic potential of 25 kcal mol–1 Å–2; hydrogens were not re-strained) using the OPLS4 force filed; water molecules > 5 Å from ligand atoms were deleted. The receptor grid was also generated in Maestro using default options. Briefly, van der Waals radius scaling factor was set to 1.0 and partial charge cutoff to 0.25; the docking box was limited to a 10 Å cube defined around the centroid of the ligand; grid-based constraint used were: hydrogen bond (main chain NH of Val270 and side chain NH of Lys194 and positional (defined as a 5 Å sphere around the ligand furopyridine moiety). The ligand (SGC-CAMKK2-1) was prepared for docking using LigPrep within Maestro using default settings. Briefly, the OPLS4 force filed was used and possible protonation states were generated at pH 7.0 ± 2.0 using Epik. Docking of SGC-CAMKK2-1 onto CAMKK2 coordinates was performed using Glide within Maestro[55] using default settings. Briefly, a van der Waals radii scaling factor of 0.8 and a partial charge cutoff of 0.15 was used; all three grid-based constrains (2x hydrogen bonds and 1x positional; as described above were enforced; docking was performed at extra precision (XP mode) with flexible ligand sampling (including nitrogen inversion and ring conformations) and Epik state penalties were added to the final docking score (−13.7565).

### 2.6 Western Blot Analysis

#### 2.6.1 Prostate Cancer Cellular Screening

C4-2 cells were plated in 6-well plates in IMEM medium containing 0.5% FBS. After 72 hours, the cells were then treated with the compounds for 24 hours before the media was aspirated and cells were washed twice in ice cold PBS. Cells were lysed using RIPA buffer containing phosphatase and protease inhibitor cocktail while rotating for 30 minutes at 4°C. In each lane, 30 □g/well of protein lysate was loaded into a 10% SDS-PAGE gel and run for 1 hour and 30 minutes. Gels were then transferred overnight in a TRIS-glycine/methanol transfer buffer onto a PVDF membrane at 4°C. Membranes were blocked, incubated with primary overnight at 4°C, washed, incubated with secondary at room temperature for 1 hour, washed, and then developed on an Azure Biosystems C-600 imager. Primary antibodies used were from Cell Signaling (Danvers, MA, USA; Phospho-AMPKα (Thr172) (40H9) Rabbit mAb: Cat#: 2535; AMPKα (D5A2) Rabbit mAb Cat#: 5831), BD Bioscience (Franklin Lakes, NJ, USA; CAMKK2 mouse mAb Cat# 610544), and Sigma (St. Louis, MO, USA; GAPDH rabbit pAb: Cat# G9545). Secondary antibody (Goat Anti-Rabbit IgG (H + L)-HRP Conjugate; Cat#:1706515) was from Bio-Rad Laboratories (Hercules, CA, USA).

#### 2.6.2 Breast Cancer Cellular Screening

MDA-MB-231 cells were treated with increasing doses of STO-609 (control), 5 and 7 (negative compound) for 24 hours as indicated. The cells were then washed three times with 2 ml of ice-cold PBS and lysed with 0.15 ml of phospho-RIPA lysis buffer (Tris-Cl pH 7.5, 50 mM; NaCl, 150 mM; NP-40, 1%; sodium deoxycholate, 0.5%; SDS, 0.05%; EDTA, 5 mM; sodium fluoride, 50 mM; sodium pyrophosphate, 15mM; ß-glyc-erophosphate, 10 mM; sodium orthovanadate, 1 mM) with protease inhibitor cocktail (Millipore-Sigma, P-8340). Equal amounts of protein per sample/lane were denatured and resolved by SDS-PAGE. Proteins were transferred to Odyssey Nitrocellulose Membranes (LI-COR Biosciences, cat no: 926-31092), and quantitative immunoblotting was performed using the Odyssey infrared immunoblotting detection system (LI-COR Biosciences, Lincoln, NE). Primary antibodies used were anti-phospho AMPKα (Thr172) (Cell Signaling, cat no: 2535, dilution 1:1000); anti-AMPKα (Cell Signaling, cat.no: 2532, dil 1:500) and anti β-actin (Cat no: 3700, Cell signaling, dil 1:10000). Secondary antibodies used were CF680 goat anti-mouse IgG (Biotium, cat no: 20065; dilution 1:15000) and CF770 goat anti-rabbit IgG (Biotium, cat no: 20078; dilution 1:15000). All antibodies were used according to the manufacturer’s instructions.

### 2.7 Pharmacokinetic Analysis

#### 2.7.1 Animals

CD-1 mice with weights 23-25 g were purchased from Charles River Laboratories.

#### 2.7.2 Animal Experiments

The animal experiments were performed at Alliance Pharma (Malvern, PA) in accordance with the guide for the care and use of laboratory animals. Alliance Pharma is accredited by the Association for Assessment and Accreditation of Laboratory Animal Care (AAALAC).

#### 2.7.3 Pharmacokinetic (PK) study of Compound 5 and STO-609 in mice

The intraperitoneal (IP) pharmacokinetics of 5 and STO-609 were evaluated by administering the compounds at a dose of 10 mg/kg via IP administration to three animals for each compound. The test compounds were dissolved in DMSO and then diluted to the appropriate concentration with the vehicle. For formulation we targeted the minimal amount of DMSO that could be used in combination with a formulation composed of 0.5% HPMC/0.2% Polysorbate 80. The final formulation vehicle for probe 5 was 5% DMSO in 0.5% HPMC/0.2% Polysorbate 80 and for STO-609 was 20% DMSO in 0.5% HPMC/0.2% Polysorbate 80. Blood was collected at t = 0.5, 1, 3, and 8 hours and immediately spun for 3 minutes and the plasma was collected, and compound levels were quantified with HPLC/MS. The raw data is available in the supplemental information.

## 3. Results

### 3.1. CAMKK2 Probe Development Synthesis and Profiling

To address the lack of a highly selective CAMKK2 inhibitor and provide the community with an additional tool to cross-validate work done using STO-609, we initiated a medicinal chemistry campaign to identify and fully characterize a CAMKK2 chemical probe. We used compound 2 (Figure 1) from Price et al.[47] as our chemical starting point, reasoning that we could improve the selectivity of this very potent starting point (pIC50 = 9.2). We hypothesized that modifying the pyrrolo[2,3-b]pyridine hinge-binding scaffold, common to many kinase inhibitors, would provide the best opportunity to increase selectivity. Our strategy to identify scaffolds with improved selectivity has been reported [52]. In short, modification of the hinge binder from the pyrrolo[2,3-b]pyridine to a furo[2,3-b]pyridine core provided the boost of selectivity needed. Furo-pyridine cores are less prevalent amongst literature kinase inhibitors, perhaps because they provide one less interaction with the hinge region of the kinase. We opted to maintain the benzoic acid that made a key interaction with the catalytic lysine in hopes that would aid in maintaining potency. The route we employed (Scheme 1) started with the 5-chlorofuro[2,3-b]pyridin-3-triflate (8). This route had been previously optimized by our group and provided a furopyridine core that could be quickly modified at both the 3 and 5-positions.[56] Compound 8 then underwent a Suzuki-Miyara reaction with 9 to preferentially install the cyclopentylbenzoic acid methyl ester at the 3-position (10). Having the chloro handle at the 5-position, we then performed a subsequent Suzuki-Miyara reaction with an aryl boronic acid to obtain the desired intermediates. Hydrolysis of the methyl ester to the carboxylic acid proceeded as expected to provide final compounds (5 and 6). The synthesis of the pinacol borane 9 was previously described [56].

**Scheme 1.**
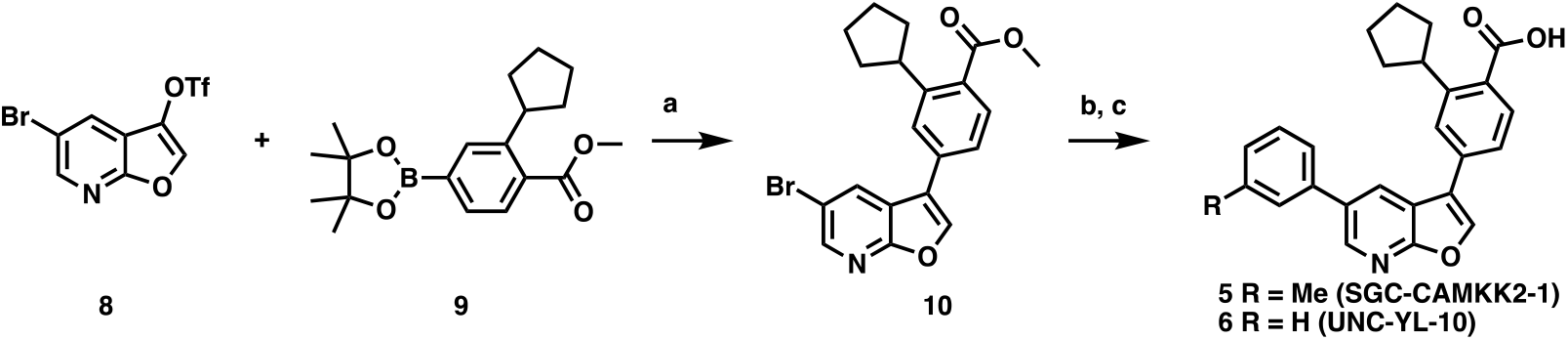
Synthetic route to compounds **5** and **6**. Reagents and conditions (a) **8**, Pd(dppf)Cl_2_ (0.05 eq.), Cs_2_CO_3_ (2 eq.), 1,4-dioxane/water (4:1), 120 °C microwave, 30 min, (b) ArB(OH)_2_ (1 eq.), Pd(dppf)Cl_2_ (0.05 eq.), Cs_2_CO_3_ (2 eq.), 1,4-dioxane/water (4:1), 120 °C microwave, 30 min, (c) 1 N NaOH, MeOH, 75 °C then aq. HCl, 1 h.

A series of furo[2,3-b]pyridines were generated with modifications off the core at the 3 and 5-positions. They were profiled in the CAMKK2 enzyme assay we have described and utilized previously. Briefly, the enzyme assay measures the ability of the compound to inhibit the transfer of a radiolabeled phosphate from ATP to a CAMKK2-specific peptide. One compound in the series, compound **5** (SGC-CAMKK2-1), although less potent than azaindole GSK650394, retained an acceptable potency (IC_50_ = 30 nM). **5** was then profiled in the DiscoverX KINOME*_scan_*™ assay panel to provide a view of the overall kinome-wide selectivity. This assay platform evaluates compound affinity for a panel of more than 400 wild-type human kinases. The S_10_ (1 μM), a calculated score that is reflective of broad kinome selectivity, is 0.002 for the probe molecule. In this panel, compound **5** only demonstrated significant affinity for CAMKK1 and CAMKK2 (CAMKK2 PoC = 5.4, CAMKK1 PoC = 12) at a compound treatment of 1 μM (Figure 2). The affinity for the next best kinase, MYO3B, is significantly lower at only 45% of control.

**Figure 2.**
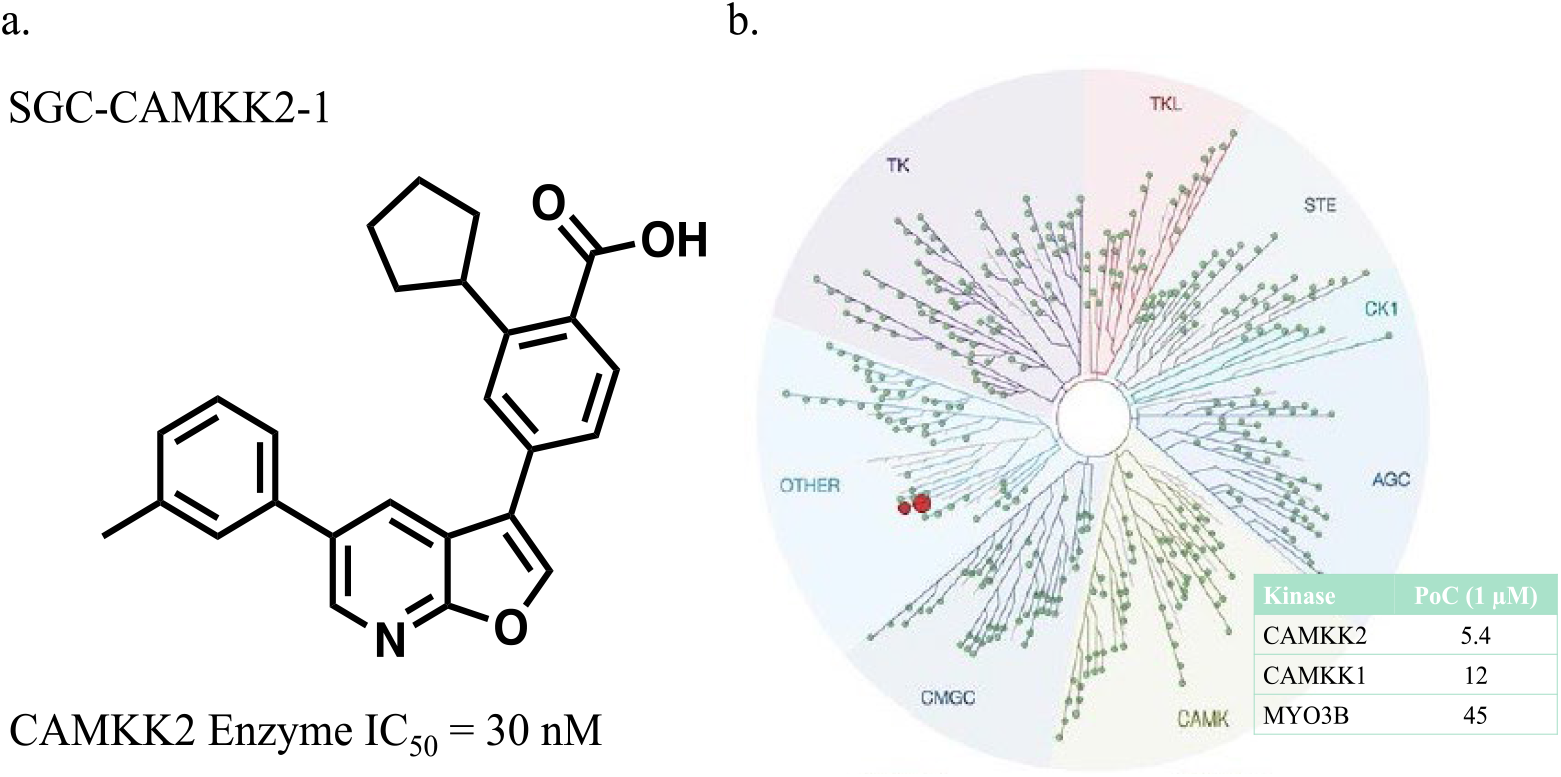
(a) The structure and enzyme inhibition IC50 of the CAMKK2 probe molecule, (b) The selectivity of compound **5** (SGC-CAMKK2-1) depicted using a TREEspot diagram obtained from profiling SGC-CAMKK2-1 at 1 μM in the KinomeSCAN^®^ assay panel.

### 3.2. Development of a structurally related negative control

We next sought to develop a structurally related but CAMKK2-inactive compound to serve as a negative control [57]. Two minor modifications to the probe (5), removal of the methyl group from the 5-position aryl ring and replacement of the cyclopentyl moiety with a chloro resulted in negative control compound 7 (SGC-CAMKK2-1N). These changes rendered this compound significantly less potent with CAMKK2 IC50 = 27 μM (Figure 3). In the DiscoverX KinomeSCAN^®^ assay panel, when screened at a concentration of 1 μM, compound 7 bound to very few kinases with S10 (1 μM) = 0.005. The only two kinases with significant binding for compound 7 were PIP4K2C with a PoC = 7.2 and PIM2 with PoC = 7.4. PIP4K2C is a lipid kinase that has very little catalytic activity[58–60], and there is no commercially available kinase activity assay, so this potential off-target activity for the negative control remains uncharacterized. We evaluated the PIM2 inhibition of negative control compound 7 at Reaction Biology Corporation and it had an IC50 = 1.9 μM.

**Figure 3.**
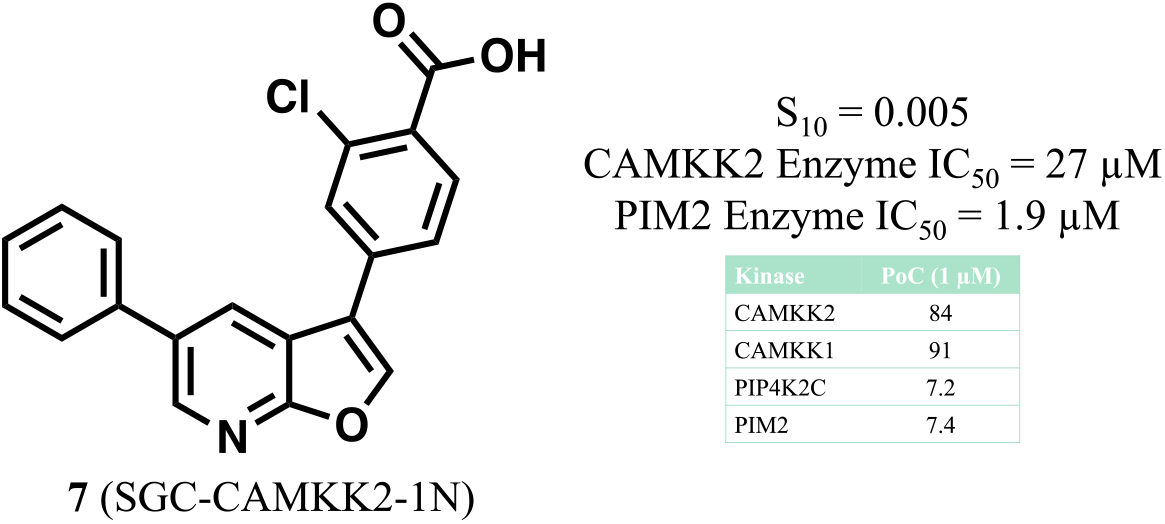
Structure and characterization of the negative control compound SGC-CAMKK2-1N.

The synthetic route to access the negative control commenced with a Suzuki-Miyaura reaction between commercially available material 5-bromofuro[2,3-b]pyridine 13 and phenylboronic acid to yield the aryl-substituted derivative 14, which was treated with bromine to afford 15 [61]. Reaction of 15 with 4-borono-2-chlorobenzoic acid afforded the carboxylic acid 7.

**Scheme 2.**
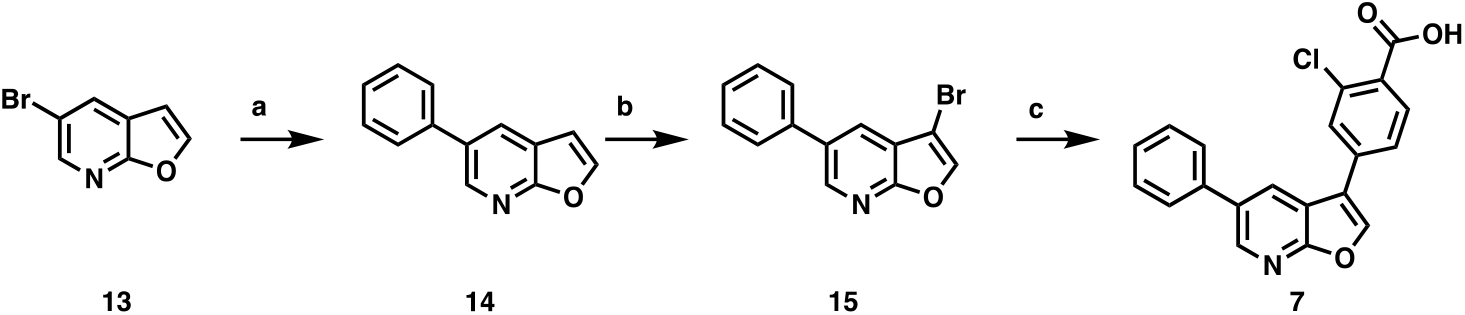
Synthesis of negative control 7. Reagents and conditions: a) PhB(OH)2, Cs2CO3, Pd(PPh3)4, DMF/H2O, 16 h; b) i. CHCl3, Br2, ii. 10% NaOH, MeOH; c) 4-borono-2-chlorobenzoic acid, Pd(PPh3)4, Na2CO3, dioxane/H2O, 90 °C, 16 h.

### 3.2. Molecular basis for ligand binding

We have previously published the crystal structure of compound 6, a close analog of the chemical probe SGC-CAMKK2-1, bound to the CAMKK2 kinase domain (KD) (amino acids 161-449) [52]. The overall structure of the complex is depicted in Figure 4. Figure 4A highlights the detailed interactions of the close analog compound 6, and figure 4B shows the predicted binding mode of the probe SGC-CAMKK2-1. The two compounds (the probe and compound 6) differ only by the presence of a methyl group in the meta position of the aryl group attached to the pyridine ring in the probe SGC-CAMKK2-1. The co-crystal structure of compound 6 bound to CAMKK2 KD revealed the ligand carboxylate moiety participates in an extensive hydrogen bond network with residues from catalytically important regions of the kinase domain, such as Asp330 in the DFG motif, Glu263 in α-helix C and Lys194. Some of these interactions are mediated by water molecules. The ligand makes an additional hydrogen bond to the main chain N atom of Val270 in the kinase domain hinge region. Given the structural similarities between the two compounds, we expect a near identical binding mode for the probe. Indeed, in silico docking of SGC-CAMKK2-1 onto the structure of CAMKK2 KD bound to 6 indicated the kinase binds to both ligands in a similar manner. In the predicted binding pose, the additional methyl group in SGC-CAMKK2-1 is sandwiched between the side chain of Pro274 and the main chain of Gly172. This latter residue is part of the P-Loop region in the kinase domain (for clarity the P-Loop is not shown in Figure 4).

**Figure 4.**
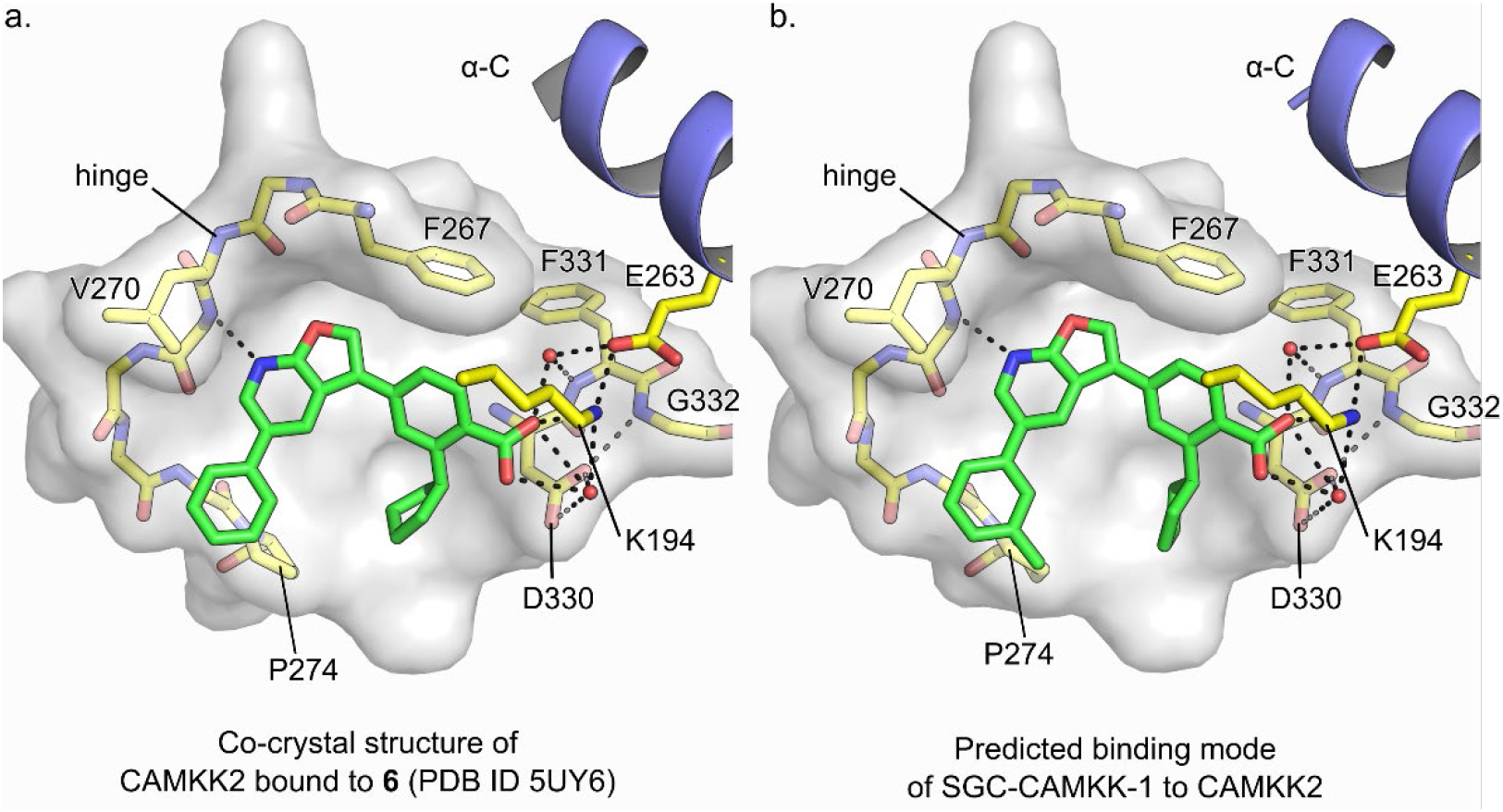
Predicted binding mode of probe SGC-CAMKK1 to CAMKK2 kinase domain. (a) Detailed view of the binding interactions between compound 6, a close furopyridine analogue of probe SGC-CAMKK-1, and CAMKK2 ATP-binding pocket as seen in the complex co-crystal structure (PDB ID 5UY6). (b) Predicted binding mode of probe SGC-CAMKK1 to CAMKK2 ATP-binding pocket obtained by in silico docking (see Methods for details). In a and b, dashed black lines indicate potential hydrogen bonds, water molecules are depicted as red spheres, carbon atoms are shown in green (ligand) or yellow (protein), the kinase domain α-helix C is shown in blue, and protein surface (for the bottom of the ATP-binding site) is shown in white.

Our binding and structural data suggest that the selectivity of 6 is a function of its single hinge contact and its ability to mediate both hydrophobic and polar interactions with CAMKK2’s ATP-binding site. Most ATP-competitive inhibitors engage the kinase hinge region via two or more hydrogen bonds to protein main chain atoms. Reducing the number of contacts between the inhibitor and the protein has been suggested as a strategy to increase selectivity [62,63].

### 3.3 CAMKK2 NanoBRET in Cell Target Engagement

In addition to in vitro enzymatic potency and kinome selectivity, a chemical probe needs to have suitable cellular potency. To evaluate this, we developed a NanoBRET in-cell target engagement assay. For this assay, the CAMKK2 kinase domain fused with a Nanoluciferase (NLuc) tag is transiently transfected into HEK293 cells. After 24 hours, a heterobifunctional molecule that consists of an inhibitor that binds to CAMKK2 linked to a BODIPY dye is added to the cells. When the inhibitor-dye hybrid binds to the kinase, the proximity to the NLuc fused to CAMKK2 creates a BRET signal. This signal can then be competed away in a dose-dependent fashion with free inhibitor. Using this assay, we evaluated the probe and STO-609 (Figure 5). SGC-CAMKK2-1 (5) was able to compete away the tracer with an in-cell target engagement IC50 of 370 nM. In the same assay, STO-609 had an IC50 > 10 μM. These data demonstrate that the probe compound can engage CAMKK2 in cells at concentrations below 1 μM.

**Figure 5.**
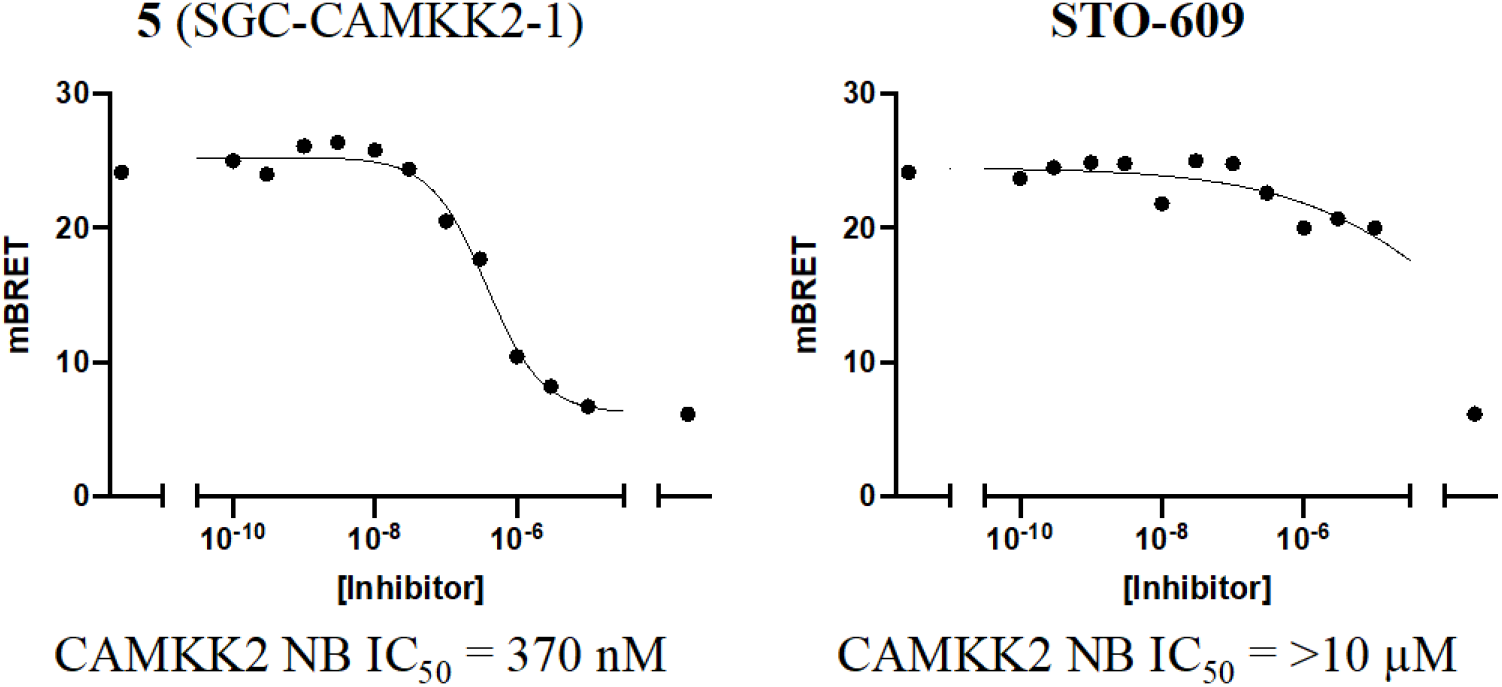
NanoBRET in-cell target engagement of CAMKK2 probe (5) and STO-609 in HEK293 cells transiently transfected with NLuc-CAMKK2. Increasing doses of inhibitor displaces the fluorescent inhibitor ligand (called a tracer) used in the assay, leading to a reduction in the BRET signal. All data was collected in n=3.

### 3.4 On-target cellular effect (western blot)

With a potent (enzyme assay) and selective (KINOMEscan) compound in hand, along with evidence of incell target engagement (nanoBRET assay), we next looked for functional consequences of CAMKK2 inhibition in cellular contexts. To do this, we evaluated the impact of our probe, the negative control, and a positive control (STO-609) on AMPK phosphorylation in C4-2 prostate cancer cells to provide evidence of a functional on-target effect. As AMPK is a direct substrate of CAMKK2, inhibition of AMPK phosphorylation at Thrl72 should be observed with the CAMKK2 chemical probe and STO-609, but not the negative control. In this experiment, IC50 was calculated via quantifying changes in p-AMPK(Thr172) normalized to total AMPK levels. The chemical probe indeed decreased p-AMPK levels, with an IC50 = 1.6 μM. As a comparison, the canonical CAMKK2 inhibitor, STO-609, also decreased p-AMPK, but was roughly 7-fold weaker with an IC50 = 10.7 μM. Importantly, the structurally similar negative control (7) did not have any measurable activity (Figure 6C) in this assay.

**Figure 6.**
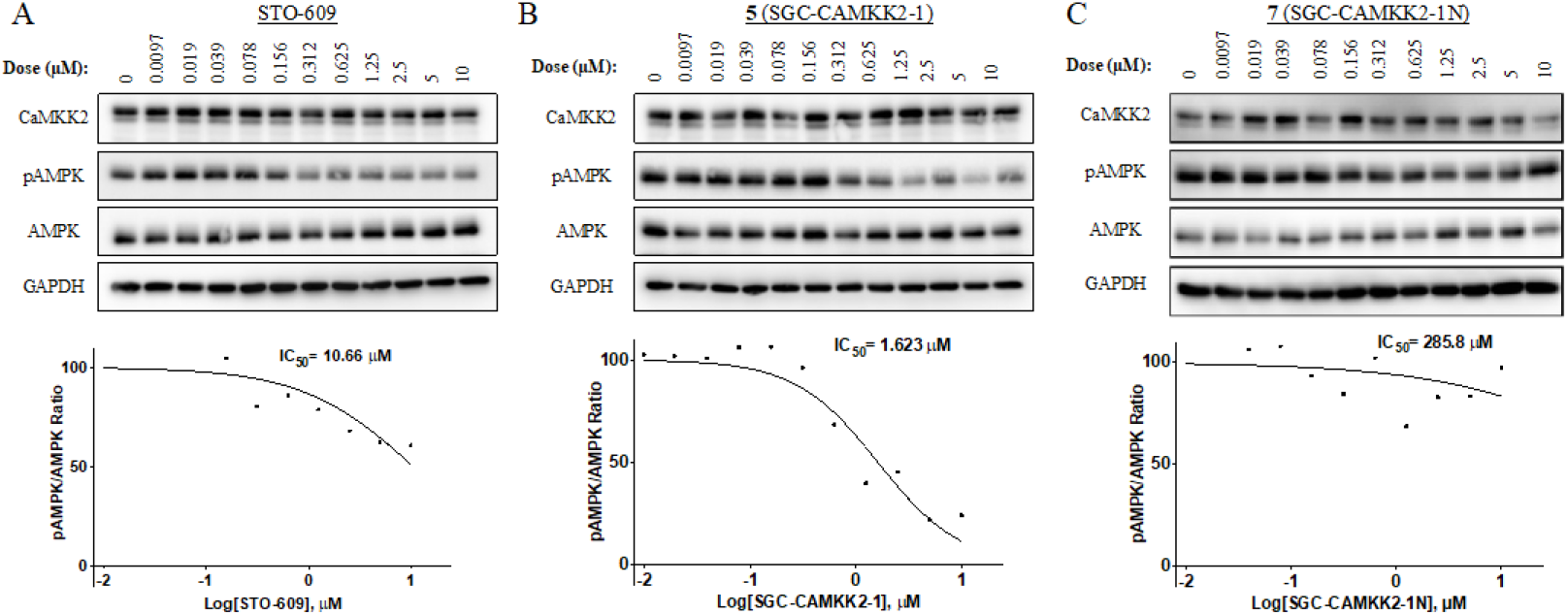
Western blot analysis of endogenous CAMKK2 inhibition using p-AMPK(Thr172) levels as a marker of CAMKK2 cellular activity. Western blot images (top) and densitometry analyses (bottom) of C4-2 cells treated with increasing doses of (A) STO-609, (B) probe SGC-CAMKK2-1 (5), and (C) negative control SGC-CAMKK2-1N (7). IC50 for STO-609 = 10.7 μM, SGC-CAMKK2-1 probe = 1.6 μM, and negative control SGC-CAMKK2-1N = >10 μM. Results represent values from three individual experiments. To confirm our cellular effects in a different model, we treated MDA-MB-231 triple-negative metastatic breast cancer cells with STO-609, probe SGC-CAMKK2-1, and negative control SGC-CAMKK2-1N. Again, SGC-CAMKK2-1 was more effective than STO-609 in blocking AMPK phosphorylation, and the negative control showed no effect at a concentration of 10 μM.

**Figure 7.**
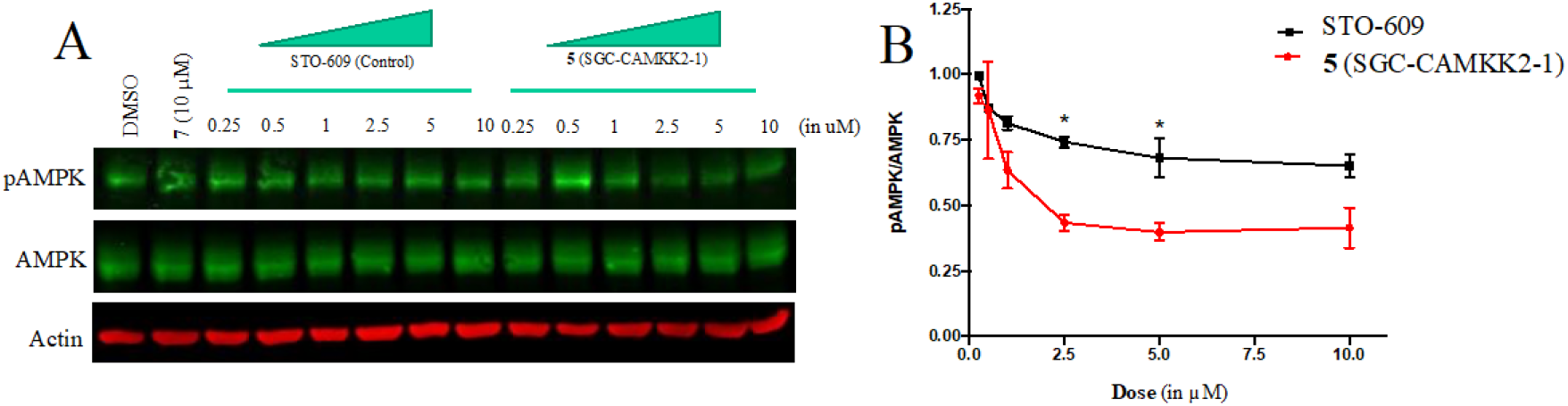
The probe compound SGC-CAMKK2-1 (5) inhibits CAMKK2 activity with higher efficacy than STO-609 in MDA-MB 231 triple-negative metastatic breast cancer cells. MDA-MB 231 cells were treated with increasing doses of STO-609 or 5 for 24 hours as indicated. 7 (10 μM) was used as the negative control for CAMKK2 inhibition (data not shown). Lysates were collected and immunoblotting was done to check for levels of p-AMPK(Thr172), a substrate of CAMKK2. A. Representative image showing 5 inhibiting CAMKK2 downstream activity with improved efficacy compared to STO-609, B. Quantification of immunoblot bands. Results represent values from three individual experiments. *P<0.05.

### 3.5 Pharmacokinetic Studies

With the probe molecule having favorable selectivity and potency in vitro and in a cellular context, we next wanted to determine its suitability for in vivo experiments. SGC-CAMKK2-1 (5) was progressed to a single dose mouse intraperitoneal (I.P) experiment. The mice were treated with either the probe or STO-609 at a concentration of 10 mg/kg.

Figure 8 shows the plasma concentration of both SGC-CAMKK2-1 and STO-609 over time. STO-609 has considerably higher plasma concentrations than SGC-CAMKK2-1 in this experiment. Although it may be possible to find a dosing regimen that allows for evaluation of this compound in vivo, based on these results we suggest only using SGC-CAMKK2-1 in cell-based assays. Further medicinal chemistry work will be needed to optimize for in vivo use.

**Figure 8.**
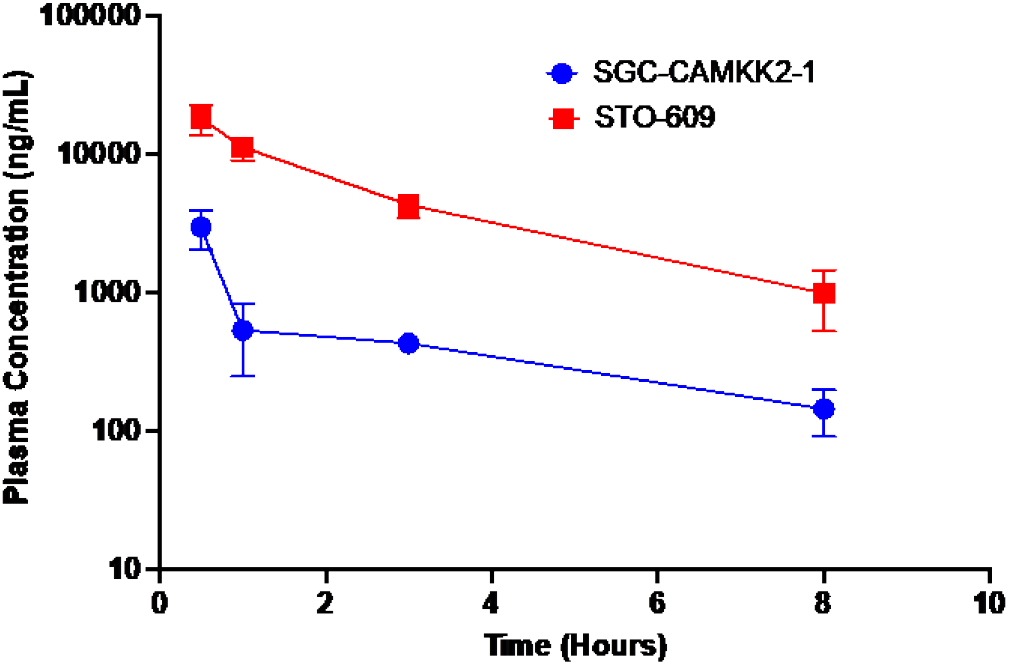
Plasma concentration of SGC-CAMKK2-1 and STO-609 in a single dose IP mouse study. The compounds were administered at a dose of 10 mg/kg via IP administration to three animals for each compound. Blood was collected at t = 0.5, 1, 3, and 8 hours and immediately spun for 3 minutes and the plasma was collected, and compound levels were quantified with HPLC/MS.

## 4. Discussion

A growing body of literature indicates both increased roles for CAMKK2 in a range of biological processes and a corresponding interest in the field. For example, CAMKK2 is a promising therapeutic target in a diverse set of diseases[29,40,64,65], including multiple cancers[7,15,19,66–69]. To date, STO-609 has been the primary chemical tool used to explore the consequences of CAMKK2 inhibition in models of disease. Although it has demonstrable on-target CAMKK2 activity and has had utility in understanding CAMKK2 roles in signaling, these studies remain limited by virtue of the observed off-target activities of STO-609[45]. Here, we describe a new chemical probe, SGC-CAMKK2-1, that can be used as an additional tool to explore CAMKK2 functions in health and disease. Advantages for SGC-CAMKK2-1 include its improved kinome selectivity and potency in cells. It has weaknesses as well, including inability to distinguish between CAMKK1 and CAMKK2 (just like STO-609), and poor *in vivo* PK in preliminary experiments. Further work should focus on identifying analogs with suitable PK properties while maintaining or improving in-cell potency and selectivity. In addition, it will be important to understand how well this compound (and STO-609) inhibit the phosphorylation of other CAMKK2 substrates such as CAMKI and CAMKIV.

SGC-CAMKK2-1 and SGC-CAMKK2-1N are commercially available through Sigma-Aldrich. As such, and with the availability of STO-609, we believe this tool kit of compounds will facilitate building a greater understanding of the true therapeutic potential of CAMKK2 inhibition. The availability of a new, readily available CAMKK2 chemical probe represents an important next step towards the ultimate goal of developing a clinical-grade CAMKK2 inhibitor to evaluate the hypothesis that inhibition of CAMKK2 will be beneficial in certain clinical settings.

Supplemental 1: Experimental details for synthesis and characterization of SGC-CAMKK2-1 and SGC-CAMKK2-1N

Supplemental 2: KINOMEscan data for the CAMKK2 probe and negative control.

## Supporting information

Supplemental 1: Experimental details for synthesis and characterization of SGC-CAMKK2-1 and SGC-CAMKK2-1N

Supplemental 2: KINOMEscan data for the CAMKK2 probe and negative control

## Author Contributions

Conceptualization, D.H.D, D.E.F, J.W.S, D.P.M, A.M, B.Y, L.R; methodology, D.H.D., C.I.W, T.L.P., C.L, D.A, J.W.S., D.E.F; validation, D.H.D, C.I.W, D. E.F, J.W.S; formal analysis, C.I.W, D.H.D, J.W.S; D.E.F; investigation, C.I.W, Y.L, T.L.P, C.L, D.A, B.E, S.O’B, M.A.H, O.G, C.G, C. C-P, P.Z, R.M.C, B.S, C.G.L, K.N, D.M, J.W.S; resources, R.M.C, B.Y, D.P.M, J.W.S, D.E.F, D.H.D; data curation, C.I.W, M.A.H., C.L., D.E.F, D.M, D.P.M, J.W.S; writing—original draft preparation, C.I.W, D. H.D, R.M.C, D.E.F, M.A.H; writing—review and editing, C.I.W, T.L.P, M.A.H, R.M.C, J.W.S, D.E.F, D.H.D; visualization, C.I.W, T.L.P., C.L, D.A, M.A.H; supervision, O.G, R.M.C, L.R, A.M, B.Y, D.P.M, J.W.S, D.E.F, D.H.D; project administration, D.H.D; funding acquisition, D.H.D, D.E.F; All authors have read and agreed to the published version of the manuscript.

## Funding

This research was funded in part by the National Cancer Institute of the National Institutes of Health (grant number R01CA218442). and was supported by the NIH Illuminating the Druggable Genome program (grant number 1U24DK116204-01). This research was also funded in part by a grant from the National Institutes of Health (NIH R01CA184208 (D.E.F.)) and a grant from the Mike Slive Foundation for Prostate Cancer Research (D.E.F.). We also acknowledge funding from FAPESP (Fundação de Amparo à Pesquisa do Estado de São Paulo) (2013/50724-5 and 2014/50897-0) and CNPq (Conselho Nacional de Desenvolvimento Científico e Tecnológico) (465651/2014-3) The SGC is a registered charity (number 1097737) that receives funds from AbbVie, Bayer Pharma AG, Boehringer Ingelheim, Canada Foundation for Innovation, Eshelman Institute for Innovation, Genome Canada, Innovative Medicines Initiative (EU/EFPIA) [ULTRA-DD grant no. 115766], Janssen, Merck KGaA Darmstadt Germany, MSD, Novartis Pharma AG, Ontario Ministry of Economic Development and Innovation, Pfizer, Takeda, and Wellcome [106169/ZZ14/Z].

## Data Availability Statement

Data describing the characterization of the chemical probe and its negative control are provide in the supplemental information. Broad kinome screening data is also included in the supplemental section.

## Acknowledgments

We are grateful for LC-MS/HRMS support provided by Dr. Brandie Ehrmann and Diane E. Wallace in the Mass Spectrometry Core Laboratory at the University of North Carolina at Chapel Hill. The core is supported by the National Science Foundation under grant no. CHE-1726291.

## Conflicts of Interest

D.E.F. has received research funding from GTx, Inc. and has familial relationships with Hummingbird Bioscience, Maia Biotechnology, Alms Therapeutics, Hinova Pharmaceuticals, and Barricade Therapeutics. The funders had no role in the conceptualization of the study or writing of the manuscript, or in the decision to publish this article.

## Notes

### Competing Interest Statement

D.E.F. has received research funding from GTx, Inc. and has familial relationships with Hummingbird Bio-science, Maia Biotechnology, Alms Therapeutics, Hinova Pharmaceuticals, and Barricade Therapeutics. The funders had no role in the conceptualization of the study or writing of the manuscript, or in the decision to publish this article.

## References

1. Anderson, K.A.; Ribar, T.J.; Lin, F.; Noeldner, P.K.; Green, M.F.; Muehlbauer, M.J.; Witters, L.A.; Kemp, B.E.; Means, A.R. Hypothalamic CaMKK2 contributes to the regulation of energy balance. Cell Metab 2008, 7, 377–388, doi:10.1016/j.cmet.2008.02.011.

2. Pritchard, Z.J.; Cary, R.L.; Yang, C.; Novack, D.V.; Voor, M.J.; Sankar, U. Inhibition of CaMKK2 reverses age-associated decline in bone mass. Bone 2015, 75, 120–127, doi:10.1016/j.bone.2015.01.021.

3. Cary, R.L.; Waddell, S.; Racioppi, L.; Long, F.; Novack, D.V.; Voor, M.J.; Sankar, U. Inhibition of Ca(2)(+)/calmodulin-dependent protein kinase kinase 2 stimulates osteoblast formation and inhibits osteoclast differentiation. J Bone Miner Res 2013, 28, 1599–1610, doi:10.1002/jbmr.1890.

4. Williams, J.N.; Kambrath, A.V.; Patel, R.B.; Kang, K.S.; Mevel, E.; Li, Y.; Cheng, Y.H.; Pucylowski, A.J.; Hassert, M.A.; Voor, M.J.; Kacena, M.A.; Thompson, W.R.; Warden, S.J.; Burr, D.B.; Allen, M.R.; Robling, A.G.; Sankar, U. Inhibition of CaMKK2 Enhances Fracture Healing by Stimulating Indian Hedgehog Signaling and Accelerating Endochondral Ossification. J Bone Miner Res 2018, 33, 930–944, doi:10.1002/jbmr.3379.

5. Broxmeyer, H.E.; Ropa, J.; Capitano, M.L.; Cooper, S.; Racioppi, L.; Sankar, U. CaMKK2 Knockout Bone Marrow Cells Collected/Processed in Low Oxygen (Physioxia) Suggests CaMKK2 as a Hematopoietic Stem to Progenitor Differentiation Fate Determinant. Stem Cell Rev Rep 2022, 18, 2513–2521, doi:10.1007/s12015-021-10306-8.

6. Racioppi, L.; Lento, W.; Huang, W.; Arvai, S.; Doan, P.L.; Harris, J.R.; Marcon, F.; Nakaya, H.I.; Liu, Y.; Chao, N. Calcium/calmodulin-dependent kinase kinase 2 regulates hematopoietic stem and progenitor cell regeneration. Cell Death Dis 2017, 8, e3076, doi:10.1038/cddis.2017.474.

7. Racioppi, L.; Nelson, E.R.; Huang, W.; Mukherjee, D.; Lawrence, S.A.; Lento, W.; Masci, A.M.; Jiao, Y.; Park, S.; York, B.; Liu, Y.; Baek, A.E.; Drewry, D.H.; Zuercher, W.J.; Bertani, F.R.; Businaro, L.; Geradts, J.; Hall, A.; Means, A.R.; Chao, N.; Chang, C.Y.; McDonnell, D.P. CaMKK2 in myeloid cells is a key regulator of the immune-suppressive microenvironment in breast cancer. Nat Commun 2019, 10, 2450, doi:10.1038/s41467-019-10424-5.

8. Huang, W.; Liu, Y.; Luz, A.; Berrong, M.; Meyer, J.N.; Zou, Y.; Swann, E.; Sundaramoorthy, P.; Kang, Y.; Jauhari, S.; Lento, W.; Chao, N.; Racioppi, L. Calcium/Calmodulin Dependent Protein Kinase Kinase 2 Regulates the Expansion of Tumor-Induced Myeloid-Derived Suppressor Cells. Front Immunol 2021, 12, 754083, doi:10.3389/fimmu.2021.754083.

9. Casciano, J.C.; Perry, C.; Cohen-Nowak, A.J.; Miller, K.D.; Vande Voorde, J.; Zhang, Q.; Chalmers, S.; Sandison, M.E.; Liu, Q.; Hedley, A.; McBryan, T.; Tang, H.Y.; Gorman, N.; Beer, T.; Speicher, D.W.; Adams, P.D.; Liu, X.; Schlegel, R.; McCarron, J.G.; Wakelam, M.J.O.; Gottlieb, E.; Kossenkov, A.V.; Schug, Z.T. MYC regulates fatty acid metabolism through a multigenic program in claudin-low triple negative breast cancer. Br J Cancer 2020, 122, 868–884, doi:10.1038/s41416-019-0711-3.

10. Jauhari, S.; Volkheimer, A.D.; Barak, I.; Guadalupe, E.; Weinberg, J.B.; Chao, N.J.A.; Li, Z.; Racioppi, L. Expression and prognostic relevance of calcium calmodulin-dependent protein kinase kinase 2 (CaMKK2) in chronic lymphocytic leukemia (CLL). Journal of Clinical Oncology 2019, 37, e19002–e19002, doi:10.1200/JCO.2019.37.15_suppl.e19002.

11. Liu, D.M.; Wang, H.J.; Han, B.; Meng, X.Q.; Chen, M.H.; Yang, D.B.; Sun, Y.; Li, Y.L.; Jiang, C.L. CAMKK2, Regulated by Promoter Methylation, is a Prognostic Marker in Diffuse Gliomas. CNS Neurosci Ther 2016, 22, 518–524, doi:10.1111/cns.12531.

12. Subbannayya, Y.; Syed, N.; Barbhuiya, M.A.; Raja, R.; Marimuthu, A.; Sahasrabuddhe, N.; Pinto, S.M.; Manda, S.S.; Renuse, S.; Manju, H.C.; Zameer, M.A.; Sharma, J.; Brait, M.; Srikumar, K.; Roa, J.C.; Vijaya Kumar, M.; Kumar, K.V.; Prasad, T.S.; Ramaswamy, G.; Kumar, R.V.; Pandey, A.; Gowda, H.; Chatterjee, A. Calcium calmodulin dependent kinase kinase 2 - a novel therapeutic target for gastric adenocarcinoma. Cancer Biol Ther 2015, 16, 336–345, doi:10.4161/15384047.2014.972264.

13. Lin, F.; Marcelo, K.L.; Rajapakshe, K.; Coarfa, C.; Dean, A.; Wilganowski, N.; Robinson, H.; Sevick, E.; Bissig, K.D.; Goldie, L.C.; Means, A.R.; York, B. The camKK2/camKIV relay is an essential regulator of hepatic cancer. Hepatology 2015, 62, 505–520, doi:10.1002/hep.27832.

14. Jin, L.; Chun, J.; Pan, C.; Kumar, A.; Zhang, G.; Ha, Y.; Li, D.; Alesi, G.N.; Kang, Y.; Zhou, L.; Yu, W.M.; Magliocca, K.R.; Khuri, F.R.; Qu, C.K.; Metallo, C.; Owonikoko, T.K.; Kang, S. The PLAG1-GDH1 Axis Promotes Anoikis Resistance and Tumor Metastasis through CamKK2-AMPK Signaling in LKB1-Deficient Lung Cancer. Mol Cell 2018, 69, 87–99 e87, doi:10.1016/j.molcel.2017.11.025.

15. Gocher, A.M.; Azabdaftari, G.; Euscher, L.M.; Dai, S.; Karacosta, L.G.; Franke, T.F.; Edelman, A.M. Akt activation by Ca(2+)/calmodulin-dependent protein kinase kinase 2 (CaMKK2) in ovarian cancer cells. J Biol Chem 2017, 292, 14188–14204, doi:10.1074/jbc.M117.778464.

16. Penfold, L.; Woods, A.; Muckett, P.; Nikitin, A.Y.; Kent, T.R.; Zhang, S.; Graham, R.; Pollard, A.; Carling, D. CAMKK2 Promotes Prostate Cancer Independently of AMPK via Increased Lipogenesis. Cancer Res 2018, 78, 6747–6761, doi:10.1158/0008-5472.CAN-18-0585.

17. Lin, C.; Blessing, A.M.; Pulliam, T.L.; Shi, Y.; Wilkenfeld, S.R.; Han, J.J.; Murray, M.M.; Pham, A.H.; Duong, K.; Brun, S.N.; Shaw, R.J.; Ittmann, M.M.; Frigo, D.E. Inhibition of CAMKK2 impairs autophagy and castration-resistant prostate cancer via suppression of AMPK-ULK1 signaling. Oncogene 2021, 40, 1690–1705, doi:10.1038/s41388-021-01658-z.

18. Saxena, N.; Beraldi, E.; Fazli, L.; Somasekharan, S.P.; Adomat, H.; Zhang, F.; Molokwu, C.; Gleave, A.; Nappi, L.; Nguyen, K.; Brar, P.; Nikesitch, N.; Wang, Y.; Collins, C.; Sorensen, P.H.; Gleave, M. Androgen receptor (AR) antagonism triggers acute succinate-mediated adaptive responses to reactivate AR signaling. EMBO Mol Med 2021, 13, e13427, doi:10.15252/emmm.202013427.

19. Pulliam, T.L.; Goli, P.; Awad, D.; Lin, C.; Wilkenfeld, S.R.; Frigo, D.E. Regulation and role of CAMKK2 in prostate cancer. Nat Rev Urol 2022, 19, 367–380, doi:10.1038/s41585-022-00588-z.

20. Pulliam, T.L.; Awad, D.; Han, J.J.; Murray, M.M.; Ackroyd, J.J.; Goli, P.; Oakhill, J.S.; Scott, J.W.; Ittmann, M.M.; Frigo, D.E. Systemic Ablation of Camkk2 Impairs Metastatic Colonization and Improves Insulin Sensitivity in TRAMP Mice: Evidence for Cancer Cell-Extrinsic CAMKK2 Functions in Prostate Cancer. Cells 2022, 11, doi:10.3390/cells11121890.

21. Zhang, Y.; Recouvreux, M.V.; Jung, M.; Galenkamp, K.M.O.; Li, Y.; Zagnitko, O.; Scott, D.A.; Lowy, A.M.; Commisso, C. Macropinocytosis in Cancer-Associated Fibroblasts Is Dependent on CaMKK2/ARHGEF2 Signaling and Functions to Support Tumor and Stromal Cell Fitness. Cancer Discov 2021, 11, 1808–1825, doi:10.1158/2159-8290.CD-20-0119.

22. Najar, M.A.; Rex, D.A.B.; Modi, P.K.; Agarwal, N.; Dagamajalu, S.; Karthikkeyan, G.; Vijayakumar, M.; Chatterjee, A.; Sankar, U.; Prasad, T.S.K. A complete map of the Calcium/calmodulin-dependent protein kinase kinase 2 (CAMKK2) signaling pathway. J Cell Commun Signal 2021, 15, 283–290, doi:10.1007/s12079-020-00592-1.

23. Racioppi, L.; Noeldner, P.K.; Lin, F.; Arvai, S.; Means, A.R. Calcium/calmodulin-dependent protein kinase kinase 2 regulates macrophage-mediated inflammatory responses. J Biol Chem 2012, 287, 11579–11591, doi:10.1074/jbc.M111.336032.

24. Teng, E.C.; Racioppi, L.; Means, A.R. A cell-intrinsic role for CaMKK2 in granulocyte lineage commitment and differentiation. J Leukoc Biol 2011, 90, 897–909, doi:10.1189/jlb.0311152.

25. Saito, M.; Saito, M.; Das, B.C. Involvement of AMP-activated protein kinase in neuroinflammation and neurodegeneration in the adult and developing brain. Int J Dev Neurosci 2019, 77, 48–59, doi:10.1016/j.ijdevneu.2019.01.007.

26. Xu, Y.; Xu, Y.; Wang, Y.; Wang, Y.; He, L.; Jiang, Z.; Huang, Z.; Liao, H.; Li, J.; Saavedra, J.M.; Zhang, L.; Pang, T. Telmisartan prevention of LPS-induced microglia activation involves M2 microglia polarization via CaMKKbeta-dependent AMPK activation. Brain Behav Immun 2015, 50, 298–313, doi:10.1016/j.bbi.2015.07.015.

27. Zhang, Y.; Xu, N.; Ding, Y.; Zhang, Y.; Li, Q.; Flores, J.; Haghighiabyaneh, M.; Doycheva, D.; Tang, J.; Zhang, J.H. Chemerin suppresses neuroinflammation and improves neurological recovery via CaMKK2/AMPK/Nrf2 pathway after germinal matrix hemorrhage in neonatal rats. Brain Behav Immun 2018, 70, 179–193, doi:10.1016/j.bbi.2018.02.015.

28. Silwal, P.; Kim, J.K.; Yuk, J.M.; Jo, E.K. AMP-Activated Protein Kinase and Host Defense against Infection. Int J Mol Sci 2018, 19, 3495, doi:10.3390/ijms19113495.

29. Williams, J.N.; Sankar, U. CaMKK2 Signaling in Metabolism and Skeletal Disease: a New Axis with Therapeutic Potential. Curr Osteoporos Rep 2019, 17, 169–177, doi:10.1007/s11914-019-00518-w.

30. Lin, F.; Ribar, T.J.; Means, A.R. The Ca2+/calmodulin-dependent protein kinase kinase, CaMKK2, inhibits preadipocyte differentiation. Endocrinology 2011, 152, 3668–3679, doi:10.1210/en.2011-1107.

31. York, B.; Li, F.; Lin, F.; Marcelo, K.L.; Mao, J.; Dean, A.; Gonzales, N.; Gooden, D.; Maity, S.; Coarfa, C.; Putluri, N.; Means, A.R. Pharmacological inhibition of CaMKK2 with the selective antagonist STO-609 regresses NAFLD. Sci Rep 2017, 7, 11793, doi:10.1038/s41598-017-12139-3.

32. Wen, L.; Chen, Z.; Zhang, F.; Cui, X.; Sun, W.; Geary, G.G.; Wang, Y.; Johnson, D.A.; Zhu, Y.; Chien, S.; Shyy, J.Y. Ca2+/calmodulin-dependent protein kinase kinase beta phosphorylation of Sirtuin 1 in endothelium is atheroprotective. Proc Natl Acad Sci U S A 2013, 110, E2420–2427, doi:10.1073/pnas.1309354110.

33. Yang, J.; Yu, J.; Li, D.; Yu, S.; Ke, J.; Wang, L.; Wang, Y.; Qiu, Y.; Gao, X.; Zhang, J.; Huang, L. Store-operated calcium entry-activated autophagy protects EPC proliferation via the CAMKK2-MTOR pathway in ox-LDL exposure. Autophagy 2017, 13, 82–98, doi:10.1080/15548627.2016.1245261.

34. Ye, C.; Zhang, D.; Zhao, L.; Li, Y.; Yao, X.; Wang, H.; Zhang, S.; Liu, W.; Cao, H.; Yu, S.; Wang, Y.; Jiang, J.; Wang, H.; Li, X.; Ying, H. CaMKK2 Suppresses Muscle Regeneration through the Inhibition of Myoblast Proliferation and Differentiation. Int J Mol Sci 2016, 17, doi:10.3390/ijms17101695.

35. Mairet-Coello, G.; Courchet, J.; Pieraut, S.; Courchet, V.; Maximov, A.; Polleux, F. The CAMKK2-AMPK kinase pathway mediates the synaptotoxic effects of Abeta oligomers through Tau phosphorylation. Neuron 2013, 78, 94–108, doi:10.1016/j.neuron.2013.02.003.

36. Peters, M.; Mizuno, K.; Ris, L.; Angelo, M.; Godaux, E.; Giese, K.P. Loss of Ca2+/calmodulin kinase kinase beta affects the formation of some, but not all, types of hippocampus-dependent long-term memory. J Neurosci 2003, 23, 9752–9760, doi:10.1523/JNEUROSCI.23-30-09752.2003.

37. Mizuno, K.; Antunes-Martins, A.; Ris, L.; Peters, M.; Godaux, E.; Giese, K. Calcium/calmodulin kinase kinase β has a male-specific role in memory formation. Neuroscience 2007, 145, 393–402.

38. Scott, J.W.; Park, E.; Rodriguiz, R.M.; Oakhill, J.S.; Issa, S.; O’Brien, M.T.; Dite, T.A.; Langendorf, C.G.; Wetsel, W.C.; Means, A.R. Autophosphorylation of CaMKK2 generates autonomous activity that is disrupted by a T85S mutation linked to anxiety and bipolar disorder. Scientific reports 2015, 5, 1–10.

39. Erhardt, A.; Lucae, S.; Unschuld, P.G.; Ising, M.; Kern, N.; Salyakina, D.; Lieb, R.; Uhr, M.; Binder, E.B.; Keck, M.E.; Muller-Myhsok, B.; Holsboer, F. Association of polymorphisms in P2RX7 and CaMKKb with anxiety disorders. J Affect Disord 2007, 101, 159–168, doi:10.1016/j.jad.2006.11.016.

40. Luo, X.J.; Li, M.; Huang, L.; Steinberg, S.; Mattheisen, M.; Liang, G.; Donohoe, G.; Shi, Y.; Chen, C.; Yue, W.; Alkelai, A.; Lerer, B.; Li, Z.; Yi, Q.; Rietschel, M.; Cichon, S.; Collier, D.A.; Tosato, S.; Suvisaari, J.; Rujescu, D.; Golimbet, V.; Silagadze, T.; Durmishi, N.; Milovancevic, M.P.; Stefansson, H.; Schulze, T.G.; Nothen, M.M.; Chen, C.; Lyne, R.; Morris, D.W.; Gill, M.; Corvin, A.; Zhang, D.; Dong, Q.; Moyzis, R.K.; Stefansson, K.; Sigurdsson, E.; Hu, F.; Moo, D.S.S.C.Z.C.; Su, B.; Gan, L. Convergent lines of evidence support CAMKK2 as a schizophrenia susceptibility gene. Mol Psychiatry 2014, 19, 774–783, doi:10.1038/mp.2013.103.

41. Sayad, A.; Ranjbaran, F.; Ghafouri-Fard, S.; Arsang-Jang, S.; Taheri, M. Expression Analysis of CYFIP1 and CAMKK2 Genes in the Blood of Epileptic and Schizophrenic Patients. J Mol Neurosci 2018, 65, 336–342, doi:10.1007/s12031-018-1106-2.

42. Liu, L.; McCullough, L.; Li, J. Genetic deletion of calcium/calmodulin-dependent protein kinase kinase beta (CaMKK beta) or CaMK IV exacerbates stroke outcomes in ovariectomized (OVXed) female mice. BMC Neurosci 2014, 15, 118, doi:10.1186/s12868-014-0118-2.

43. Sun, P.; Bu, F.; Min, J.W.; Munshi, Y.; Howe, M.D.; Liu, L.; Koellhoffer, E.C.; Qi, L.; McCullough, L.D.; Li, J. Inhibition of calcium/calmodulin-dependent protein kinase kinase (CaMKK) exacerbates impairment of endothelial cell and blood-brain barrier after stroke. Eur J Neurosci 2019, 49, 27–39, doi:10.1111/ejn.14223.

44. Tokumitsu, H.; Inuzuka, H.; Ishikawa, Y.; Ikeda, M.; Saji, I.; Kobayashi, R. STO-609, a specific inhibitor of the Ca(2+)/calmodulin-dependent protein kinase kinase. J Biol Chem 2002, 277, 15813–15818, doi:10.1074/jbc.M201075200.

45. O’Byrne, S.N.; Scott, J.W.; Pilotte, J.R.; Santiago, A.D.S.; Langendorf, C.G.; Oakhill, J.S.; Eduful, B.J.; Counago, R.M.; Wells, C.I.; Zuercher, W.J.; Willson, T.M.; Drewry, D.H. In Depth Analysis of Kinase Cross Screening Data to Identify CAMKK2 Inhibitory Scaffolds. Molecules 2020, 25, doi:10.3390/molecules25020325.

46. Edwards, A.M.; Bountra, C.; Kerr, D.J.; Willson, T.M. Open access chemical and clinical probes to support drug discovery. Nature Chemical Biology 2009, 5, 436–440, doi:10.1038/nchembio0709-436.

47. Price, D.J.; Drewry, D.H.; Schaller, L.T.; Thompson, B.D.; Reid, P.R.; Maloney, P.R.; Liang, X.; Banker, P.; Buckholz, R.G.; Selley, P.K.; McDonald, O.B.; Smith, J.L.; Shearer, T.W.; Cox, R.F.; Williams, S.P.; Reid, R.A.; Tacconi, S.; Faggioni, F.; Piubelli, C.; Sartori, I.; Tessari, M.; Wang, T.Y. An orally available, brain-penetrant CAMKK2 inhibitor reduces food intake in rodent model. Bioorg Med Chem Lett 2018, 28, 1958–1963, doi:10.1016/j.bmcl.2018.03.034.

48. Fabian, M.A.; Biggs, W.H., 3rd; Treiber, D.K.; Atteridge, C.E.; Azimioara, M.D.; Benedetti, M.G.; Carter, T.A.; Ciceri, P.; Edeen, P.T.; Floyd, M.; Ford, J.M.; Galvin, M.; Gerlach, J.L.; Grotzfeld, R.M.; Herrgard, S.; Insko, D.E.; Insko, M.A.; Lai, A.G.; Lelias, J.M.; Mehta, S.A.; Milanov, Z.V.; Velasco, A.M.; Wodicka, L.M.; Patel, H.K.; Zarrinkar, P.P.; Lockhart, D.J. A small molecule-kinase interaction map for clinical kinase inhibitors. Nat Biotechnol 2005, 23, 329–336, doi:10.1038/nbt1068.

49. Wodicka, L.M.; Ciceri, P.; Davis, M.I.; Hunt, J.P.; Floyd, M.; Salerno, S.; Hua, X.H.; Ford, J.M.; Armstrong, R.C.; Zarrinkar, P.P.; Treiber, D.K. Activation state-dependent binding of small molecule kinase inhibitors: structural insights from biochemistry. Chem Biol 2010, 17, 1241–1249, doi:10.1016/j.chembiol.2010.09.010.

50. O’Brien, M.T.; Oakhill, J.S.; Ling, N.X.; Langendorf, C.G.; Hoque, A.; Dite, T.A.; Means, A.R.; Kemp, B.E.; Scott, J.W. Impact of Genetic Variation on Human CaMKK2 Regulation by Ca(2+)-Calmodulin and Multisite Phosphorylation. Sci Rep 2017, 7, 43264, doi:10.1038/srep43264.

51. Vasta, J.D.; Corona, C.R.; Wilkinson, J.; Zimprich, C.A.; Hartnett, J.R.; Ingold, M.R.; Zimmerman, K.; Machleidt, T.; Kirkland, T.A.; Huwiler, K.G.; Ohana, R.F.; Slater, M.; Otto, P.; Cong, M.; Wells, C.I.; Berger, B.T.; Hanke, T.; Glas, C.; Ding, K.; Drewry, D.H.; Huber, K.V.M.; Willson, T.M.; Knapp, S.; Muller, S.; Meisenheimer, P.L.; Fan, F.; Wood, K.V.; Robers, M.B. Quantitative, Wide-Spectrum Kinase Profiling in Live Cells for Assessing the Effect of Cellular ATP on Target Engagement. Cell Chem Biol 2018, 25, 206–214 e211, doi:10.1016/j.chembiol.2017.10.010.

52. Eduful, B.J.; O’Byrne, S.N.; Temme, L.; Asquith, C.R.M.; Liang, Y.; Picado, A.; Pilotte, J.R.; Hossain, M.A.; Wells, C.I.; Zuercher, W.J.; Catta-Preta, C.M.C.; Zonzini Ramos, P.; Santiago, A.S.; Counago, R.M.; Langendorf, C.G.; Nay, K.; Oakhill, J.S.; Pulliam, T.L.; Lin, C.; Awad, D.; Willson, T.M.; Frigo, D.E.; Scott, J.W.; Drewry, D.H. Hinge Binder Scaffold Hopping Identifies Potent Calcium/Calmodulin-Dependent Protein Kinase Kinase 2 (CAMKK2) Inhibitor Chemotypes. Journal of medicinal chemistry 2021, 64, 10849–10877, doi:10.1021/acs.jmedchem.0c02274.

53. Sastry, G.M.; Adzhigirey, M.; Day, T.; Annabhimoju, R.; Sherman, W. Protein and ligand preparation: parameters, protocols, and influence on virtual screening enrichments. J Comput Aided Mol Des 2013, 27, 221–234, doi:10.1007/s10822-013-9644-8.

54. Olsson, M.H.; Søndergaard, C.R.; Rostkowski, M.; Jensen, J.H. PROPKA3: Consistent Treatment of Internal and Surface Residues in Empirical pKa Predictions. J Chem Theory Comput 2011, 7, 525–537, doi:10.1021/ct100578z.

55. Friesner, R.A.; Murphy, R.B.; Repasky, M.P.; Frye, L.L.; Greenwood, J.R.; Halgren, T.A.; Sanschagrin, P.C.; Mainz, D.T. Extra precision glide: docking and scoring incorporating a model of hydrophobic enclosure for protein-ligand complexes. Journal of medicinal chemistry 2006, 49, 6177–6196, doi:10.1021/jm051256o.

56. O’Byrne, S.N.; Eduful, B.J.; Willson, T.M.; Drewry, D.H. Concise, gram-scale synthesis of furo[2,3-b]pyridines with functional handles for chemoselective cross-coupling. Tetrahedron Lett 2020, 61 152353, doi:10.1016/j.tetlet.2020.152353.

57. Bunnage, M.E.; Chekler, E.L.; Jones, L.H. Target validation using chemical probes. Nat Chem Biol 2013, 9, 195–199, doi:10.1038/nchembio.1197.

58. Poli, A.; Zaurito, A.E.; Abdul-Hamid, S.; Fiume, R.; Faenza, I.; Divecha, N. Phosphatidylinositol 5 Phosphate (PI5P): From Behind the Scenes to the Front (Nuclear) Stage. International Journal of Molecular Sciences 2019, 20, 2080.

59. Clarke, J.H.; Irvine, R.F. Evolutionarily conserved structural changes in phosphatidylinositol 5-phosphate 4-kinase (PI5P4K) isoforms are responsible for differences in enzyme activity and localization. Biochem J 2013, 454, 49–57, doi:10.1042/bj20130488.

60. Clarke, J.H.; Giudici, M.L.; Burke, J.E.; Williams, R.L.; Maloney, D.J.; Marugan, J.; Irvine, R.F. The function of phosphatidylinositol 5-phosphate 4-kinase γ (PI5P4Kγ) explored using a specific inhibitor that targets the PI5P-binding site. Biochem J 2015, 466, 359–367, doi:10.1042/bj20141333.

61. Saitoh, M.; Kunitomo, J.; Kimura, E.; Iwashita, H.; Uno, Y.; Onishi, T.; Uchiyama, N.; Kawamoto, T.; Tanaka, T.; Mol, C.D.; Dougan, D.R.; Textor, G.P.; Snell, G.P.; Takizawa, M.; Itoh, F.; Kori, M. 2-{3-[4-(Alkylsulfinyl)phenyl]-1-benzofuran-5-yl}-5-methyl-1,3,4-oxadiazole derivatives as novel inhibitors of glycogen synthase kinase-3beta with good brain permeability. Journal of medicinal chemistry 2009, 52, 6270–6286, doi:10.1021/jm900647e.

62. Ghose, A.K.; Herbertz, T.; Pippin, D.A.; Salvino, J.M.; Mallamo, J.P. Knowledge based prediction of ligand binding modes and rational inhibitor design for kinase drug discovery. Journal of medicinal chemistry 2008, 51, 5149–5171, doi:10.1021/jm800475y.

63. Davis, M.I.; Hunt, J.P.; Herrgard, S.; Ciceri, P.; Wodicka, L.M.; Pallares, G.; Hocker, M.; Treiber, D.K.; Zarrinkar, P.P. Comprehensive analysis of kinase inhibitor selectivity. Nat Biotechnol 2011, 29, 1046–1051, doi:10.1038/nbt.1990.

64. Wu, F.; Hill, K.; Fang, Q.; He, Z.; Zheng, H.; Wang, X.; Xiong, H.; Sha, S.H. Traumatic-noise-induced hair cell death and hearing loss is mediated by activation of CaMKKβ. Cell Mol Life Sci 2022, 79, 249, doi:10.1007/s00018-022-04268-4.

65. McArdle, J.; Schafer, X.L.; Munger, J. Inhibition of calmodulin-dependent kinase kinase blocks human cytomegalovirus-induced glycolytic activation and severely attenuates production of viral progeny. J Virol 2011, 85, 705–714, doi:10.1128/jvi.01557-10.

66. Stewart, L.M.; Gerner, L.; Rettel, M.; Stein, F.; Burrows, J.F.; Mills, I.G.; Evergren, E. CaMKK2 facilitates Golgi-associated vesicle trafficking to sustain cancer cell proliferation. Cell Death Dis 2021, 12, 1040, doi:10.1038/s41419-021-04335-x.

67. Chen, Z.; Sun, X.; Xia, Z.; Wang, J.; Guo, N.; Zhang, Y. CaMKK2 Promotes the Progression of Ovarian Carcinoma through the PI3K/PDK1/Akt Axis. Comput Math Methods Med 2022, 2022, 7187940, doi:10.1155/2022/7187940.

68. Najar, M.A.; Aravind, A.; Dagamajalu, S.; Sidransky, D.; Ashktorab, H.; Smoot, D.T.; Gowda, H.; Prasad, T.S.K.; Modi, P.K.; Chatterjee, A. Hyperactivation of MEK/ERK pathway by Ca(2+) /calmodulin-dependent protein kinase kinase 2 promotes cellular proliferation by activating cyclin-dependent kinases and minichromosome maintenance protein in gastric cancer cells. Mol Carcinog 2021, 60, 769–783, doi:10.1002/mc.23343.

69. Huang, Y.K.; Su, Y.F.; Lieu, A.S.; Loh, J.K.; Li, C.Y.; Wu, C.H.; Kuo, K.L.; Lin, C.L. MiR-1271 regulates glioblastoma cell proliferation and invasion by directly targeting the CAMKK2 gene. Neurosci Lett 2020, 737, 135289, doi:10.1016/j.neulet.2020.135289.

