## Supplemental 1: Experimental details for synthesis and characterization of SGC-CAMKK2-1 and SGC-CAMKK2-1N for "SGC-CAMKK2-1: A chemical probe for CAMKK2"

General Chemistry Information

All reagents and solvents, unless specifically stated, were used as obtained from their commercial sources without further purification. Air and moisture sensitive reactions were performed under an inert atmosphere using nitrogen in a previously oven-dried or flame-dried reaction flask, and addition of reagents were done using a syringe. All microwave (MW) reactions were carried out in a Biotage Initiator EXP US 400W microwave synthesizer. Thin layer chromatography (TLC) analyses were performed using 200 μm pre-coated sorbtech fluorescent TLC plates and spots were visualized using UV light. High resolution mass spectrometry samples were analyzed with a ThermoFisher Q Exactive HF-X (ThermoFisher, Bremen, Germany) mass spectrometer coupled with a Waters Acquity H-class liquid chromatograph system. Column chromatography was undertaken with a Biotage Isolera One instrument. Nuclear magnetic resonance (NMR) spectrometry was run on a varian Inova 400 MHz or Bruker Avance III 700 MHz spectrometer equipped with a TCI H-C/N-D 5 mm cryoprobe and data was processed using the MestReNova processor. Chemical shifts are reported in ppm with residual solvent peaks referenced as internal standard.

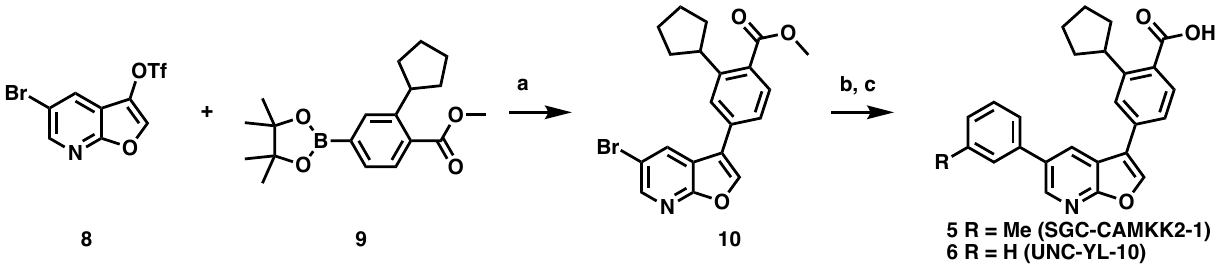

Scheme 1. Reagent and conditions: a) tetrakis (0.10 eq.), Na_2_CO_3_ (3 eq.), MeOH (2.0 mL), DCM (0.5 mL), 115 °C, 10 min, MW, 59%; b) Pd_2_(dba)_3_ (0.05 eq.), XPhos (0.01 eq.) and Cs_2_CO_3_ (3 eq.), dioxane:water (10:1, 2 mL), 120 °C, 16 h, 84%; c) 1 N NaOH, 75 °C, 4 h.

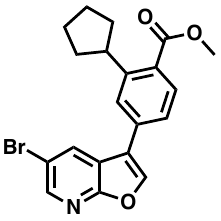

Synthesis of **methyl 4-(5-bromofuro[2,3-*b*]pyridin-3-yl)-2-cyclopentylbenzoate (10)**:

A microwave vial was charged with 5-bromofuro[2,3-b]pyridin-3-yl trifluoromethanesulfonate (250 mg, 0.72 mmol, 1 eq.) ,  methyl 2-cyclopentyl-4-(4,4,5,5-tetramethyl-1,3,2-dioxaborolan-2-yl)benzoate (358 mg, 1.08 mmol, 1 eq.), palladiumtetrakis (83.50 mg, 0.72 mmol, 0.10 eq.),  sodium carbonate (230 mg, 2.17 mmol, 3 eq.),  MeOH (2.0 mL) and  DCM (0.5 mL) . The vial was sealed and irradiated in a microwave synthesizer at 115 ^o^C for 10 min. Vial content was neutralized with aq. HCl and and extracted with ethyl acetate. The organic phases were combined and dried over sodium sulfate, concentrated and purified using flash column chromatography eluting with EtOAc/Hexanes to afford methyl 4-(5-bromofuro[2,3-b]pyridin-3-yl)-2-cyclopentylbenzoate**, 10 (59%)** was obtained as an off-white solid. ^1^H NMR (400 MHz, DMSO-d6) δ 8.74 (s, 1H), 8.56 (d, J = 1.8 Hz, 1H), 8.48 (d, J = 1.6 Hz, 1H), 7.75 (d, J = 8.2 Hz, 2H), 7.69 (dd, J = 7.9, 1.7 Hz, 1H), 3.85 (s, 3H), 3.71 – 3.58 (m, 1H), 2.08 – 1.96 (m, 2H), 1.87 – 1.76 (m, 2H), 1.73 – 1.61 (m, 4H). ^13^C NMR (101 MHz, DMSO-*d*_6_) δ 167.87, 160.49, 147.04, 144.88, 144.75, 133.42, 132.34, 130.17, 130.01, 124.98, 124.15, 119.61, 119.46, 115.29, 52.16, 41.43, 34.20, 25.19.

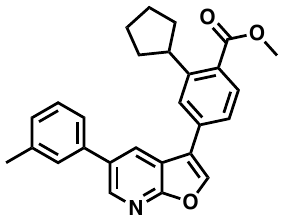

**methyl 2-cyclopentyl-4-(5-(***m***-tolyl)furo[2,3-***b***]pyridin-3-yl)benzoate:** 4-(5-bromofuro[2,3-b]pyridin-3-yl)-2-cyclopentylbenzoate (100 mg, 0.25 mmol) and 3-methylphenylboronic acid (51 mg, 0.38 mmol) were were dissolved in a 10:1 mixture of dioxane and water (2.0 mL). Pd_2_(dba)_3_ (11 mg, 0.013 mmol), XPhos (12 mg, 0.025 mmol) and cesium carbonate (244 mg, 0.75 mmol) were added. The flask was flushed with nitrogen gas and stirred at 120 ºC for 16 h. Once cooled the volatiles were removed *in vacuo*, and the crude purified by column chromatography (5% EtOAc/hexane) to afford the biaryl furopyridine (86 mg, 84%) as a white solid. 1H NMR (400 MHz, CDCl_3_) δ 8.60 (d, J = 2.2 Hz, 1H), 8.28 (d, J = 2.2 Hz, 1H), 7.98 (s, 1H), 7.88 (dd, J = 8.1, 0.4 Hz, 1H), 7.68 (d, J = 1.8 Hz, 1H), 7.50 (dd, J = 8.0, 1.8 Hz, 1H), 7.45 – 7.37 (m, 3H), 7.24 (dtd, J = 6.6, 1.8, 0.7 Hz, 1H), 3.93 (s, 3H), 3.91 – 3.81 (m, 1H), 2.46 (d, J = 0.6 Hz, 3H), 2.23 – 2.12 (m, 2H), 1.93 – 1.82 (m, 2H), 1.80 – 1.72 (m, 2H), 1.70 – 1.62 (m, 2H).

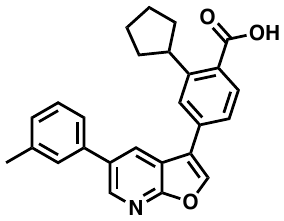

**2-cyclopentyl-4-(5-(***m***-tolyl)furo[2,3-***b***]pyridin-3-yl)benzoic acid (5):** A methyl 2-cyclopentyl-4-(5-(m-tolyl)furo[2,3-b]pyridin-3-yl)benzoate (50 mg) in methanol was treated with 1 N NaOH (aq.) and stirred for 4 hours at 75 °C. Upon completion of the reaction the reaction mixture was acidified with 2 N HCl (aq.). Pure product was extracted with ethyl acetate and crystalized from DCM/hexane. 1H NMR (400 MHz, DMSO-*d_6_*) δ 12.93 (s, 1H), 8.69 (s, 1H), 8.63 (d, J = 2.2 Hz, 1H), 8.49 (d, J = 2.2 Hz, 1H), 7.85 – 7.71 (m, 3H), 7.65 – 7.49 (m, 2H), 7.39 (t, J = 7.6 Hz, 1H), 7.29 – 7.19 (m, 1H), 3.89 – 3.73 (m, 1H), 2.39 (s, 3H), 2.14 – 1.95 (m, 2H), 1.81 (d, J = 11.5 Hz, 2H), 1.75 – 1.56 (m, 4H). ^13^C NMR (176 MHz, DMSO-*d_6_*) δ 169.73, 162.05, 147.59, 144.26, 143.69, 138.83, 137.89, 134.13, 133.46, 131.44, 130.77, 129.49, 128.94, 128.59, 125.54, 125.00, 124.60, 120.67, 118.08, 41.65, 34.86, 25.81, 21.55. HRMS m/z: [M+H]^+^ Calc’d for C_26_H_23_NO_3_ 398.17; Found 398.16.

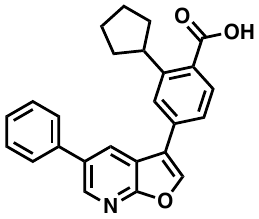

Synthesis of **2-cyclopentyl-4-(5-phenylfuro[2,3-b]pyridin-3-yl)benzoic acid**, **(6)**: A mixture of 4-(5-bromofuro[2,3-b]pyridin-3-yl)-2-cyclopentylbenzoic acid (48.0 mg, 0.12 mmol), phenylboronic acid (30.0 mg, 0.25 mmol), sodium carbonate (40.0 mg, 0.37 mmol) and Pd(PPh_3_)_4_ (14.0 mg, 12 µmol) in a mixture of Dioxane (2 mL) and Water (0.5 mL) was heated to 90 ^o^C under nitrogen overnight. Contents were neutralized with aq. HCl and extracted with ethyl acetate. The organic phases were combined, dried over anhydrous sodium sulfate, concentrated and purified using flash column chromatography eluting with EtOAc/Hexanes to obtain the target compound, 2-cyclopentyl-4-(5-phenylfuro[2,3-b]pyridin-3-yl)benzoic acid **6** (39 mg, 82 %) as a white solid. ^1^H NMR (400 MHz, DMSO-d6) δ 12.95 (s, 1H), 8.72 (s, 1H), 8.67 (d, J = 2.2 Hz, 1H), 8.53 (d, J = 2.2 Hz, 1H), 7.82 (d, J = 1.5 Hz, 1H), 7.81 – 7.79 (m, 4H), 7.56 – 7.50 (m, 2H), 7.47 – 7.41 (m, 1H), 3.86 – 3.77 (m, 1H), 2.11 – 2.01 (m, 2H), 1.87 – 1.79 (m, 2H), 1.73 – 1.62 (m, 4H). ^13^C NMR (101 MHz, DMSO-d6) δ 169.27, 161.59, 147.04, 143.81, 143.24, 137.50, 133.62, 132.88, 131.00, 130.27, 129.10, 128.18, 127.83, 127.46, 125.02, 124.13, 120.20, 117.61, 41.22, 34.35, 25.28. HRMS m/z: [M+H]^+^ Calc’d for C_25_H_21_NO_3_ 384.1600; Found 384.1593. MP Range 259-262.

Intermediate **9** was synthesized as previously described by Eduful et al.[1] Briefly **11** was treated with cylclopentylmagnesium bromide to afford intermediate **12**. The pinocol borane was then installed under standard conditions to obtain **9**.

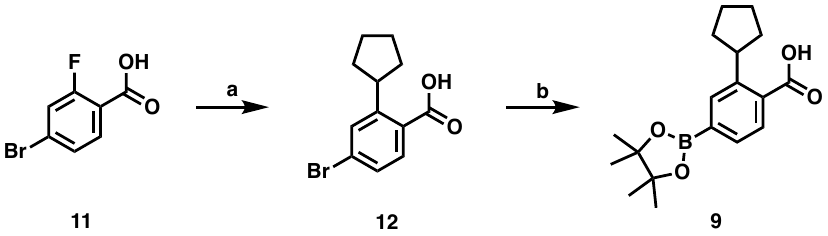

Scheme 2. Synthetic route to obtain key intermediate **9**: Reagents and conditions: a) Cyclopropylmagnesium bromide, THF, -10 to 25 °C, 5 h, then aq. HCl; b) Pd(dppf)Cl_2_.DCM (0.05 eq.), KOAc, bis(pinacolato)diboron (2 eq.), 1,4-dioxane, 100 °C, 2 h

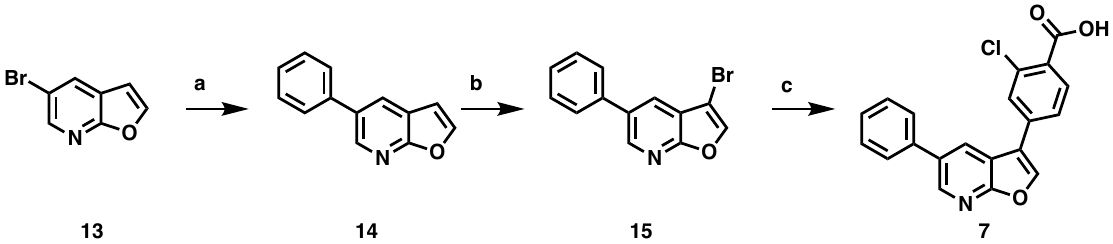

Scheme 3. Synthesis of compound 7. Reagents and conditions: a) tetrakis (0.10 eq.), Na_2_CO_3_ (3 eq.), DMF (20.0 mL), water (5.0 mL), 90 °C, 16 h, heating, 68%; b) DCM, Br_2_, 0 °C to r.t, 1 h , 44%; c) tetrakis (0.10 eq.), Na_2_CO_3_ (3 eq.), dioxane (10.0 mL), water (2.5 mL), 90 °C, 16 h, heating, 83%

**5-phenylfuro[2,3-*b*]pyridine (14):**

A mixture of 5-bromofuro[2,3-*b*]pyridine (1.00 g, 5.05 mmol), phenylboronic acid (924 mg, 7.57 mmol), palladiumtetrakis (292 mg, 0.25 mmol), and cesium carbonate (3.29 g, 10.1 mmol) in DMF (20 mL) and Water (5.0 mL) under nitrogen atmosphere was heated to 90 ^o^C for 16 h. The reaction mixture was allowed to cool to room temperature and water (100 mL) was added and extracted into EtOAc (150 mL, 2X). The combined organic layer was washed with brine (20 mL), dried over sodium sulfate, and concentrated in vacuo. The residue was purified by silica gel column chromatography (0-20% EtOAc/hexanes), to afford 5-phenylfuro[2,3-*b*]pyridine, **14** (0.67 g, 68 %) as a pale solid. LCMS [M+H]^+^: *m/z* 195.08.

**3-bromo-5-phenylfuro[2,3-b]pyridine (15):**

To a mixture of 5-phenylfuro[2,3-*b*]pyridine (630 mg, 3.23 mmol) in CHCl_3_ (25 mL) at 0 °C was slowly added bromine (183 µL, 3.55 mmol) and the mixture was stirred at room temperature for 1.5 hr. The organic layer was washed with 1 M sodium sulfite and brine, dried over sodium sulfate, and concentrated in vacuo to give 2,3-dibromo-5-phenylfuro[2,3-*b*]pyridine. To a solution of 2,3-dibromo-5-phenylfuro[2,3-*b*]pyridine in THF (20 mL) at 0 °C was added a mixture of NaOH (182 mg, 85% Wt, 3.87 mmol) in MeOH (5 mL), and the mixture was stirred for 30 min. The reaction mixture was diluted with EtOAc, and the organic layer was washed with water and saturated aqueous sodium hydrogen carbonate, dried over sodium sulfate, and concentrated in vacuo. The residue was purified by silica gel column
chromatography eluting with 0 - 15% EtOAc/hexanes to afford 3-bromo-5-phenylfuro[2,3-*b*]pyridine, **15** (390 mg, 44%) as a brownish solid. LCMS [M+H]^+^: *m/z* 272.12/274.12.

**2-chloro-4-(5-phenylfuro[2,3-b]pyridin-3-yl)benzoic acid (7):**

A mixture of 3-bromo-5-phenylfuro[2,3-*b*]pyridine (350 mg, 1.28 mmol), 4-borono-2-chlorobenzoic acid (435 mg, 2.17 mmol), palladiumtetrakis (103 mg, 89.4 µmol) and sodium carbonate (541 mg, 5.11 mmol) in dioxane (10 mL) and water (2.5 mL) under nitrogen atmosphere was heated to 90 ^o^C for 16 h. To the reaction mixture was added water (50 mL) and extracted into EtOAc (80 mL, 2x). The combined organic layer was washed with brine (20 mL), dried over sodium sulfate, and concentrated in vacuo. The residue was purified by silica gel column chromatography (45% EtOAc/hexanes), to afford 2-chloro-4-(5-phenylfuro[2,3-*b*]pyridin-3-yl)benzoic acid **7** (0.37 g, 1.1 mmol, 83 %) as a near yellow solid. ^1^H NMR (400 MHz, DMSO-*d*_6_) δ 13.43 (s, 1H), 8.79 (s, 1H), 8.66 (dd, *J* = 17.0, 2.2 Hz, 2H), 8.04 – 7.98 (m, 2H), 7.94 (d, *J* = 8.1 Hz, 1H), 7.86 – 7.82 (m, 2H), 7.56 – 7.51 (m, 2H), 7.47 – 7.42 (m, 1H). ^13^C NMR (176 MHz, DMSO-*d_6_*) δ 166.80, 162.00, 145.21, 143.95, 137.87, 135.65, 133.59, 133.18, 132.34, 130.47, 129.54, 128.95, 128.94, 128.34, 128.11, 126.04, 119.28, 117.61. HRMS calcd for C_20_H_12_ClNO_3_ [M+H]^+^: *m/z* 350.0584; found 350.0568.

^1^H NMR for **5**

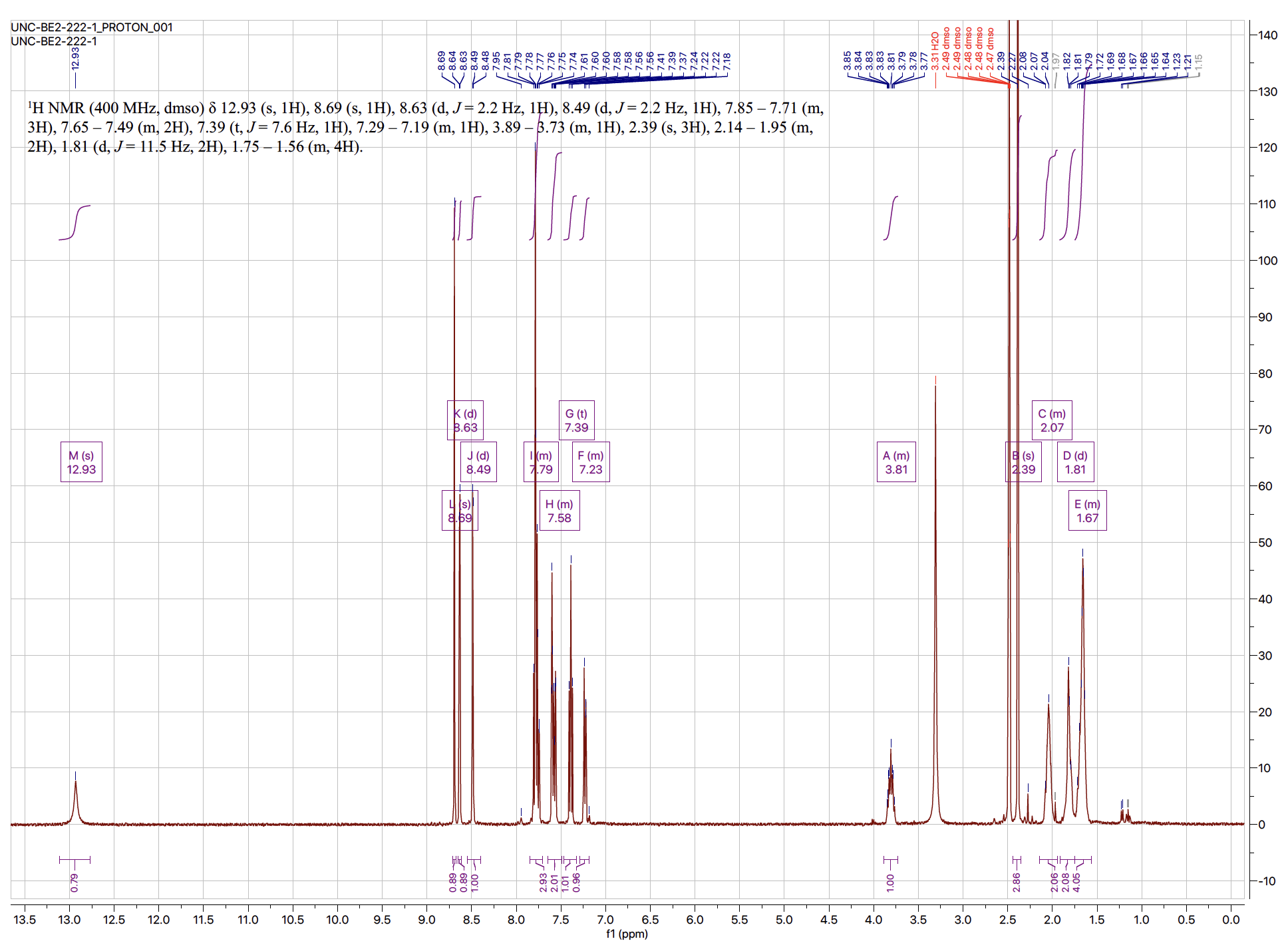

^13^C NMR for **5**

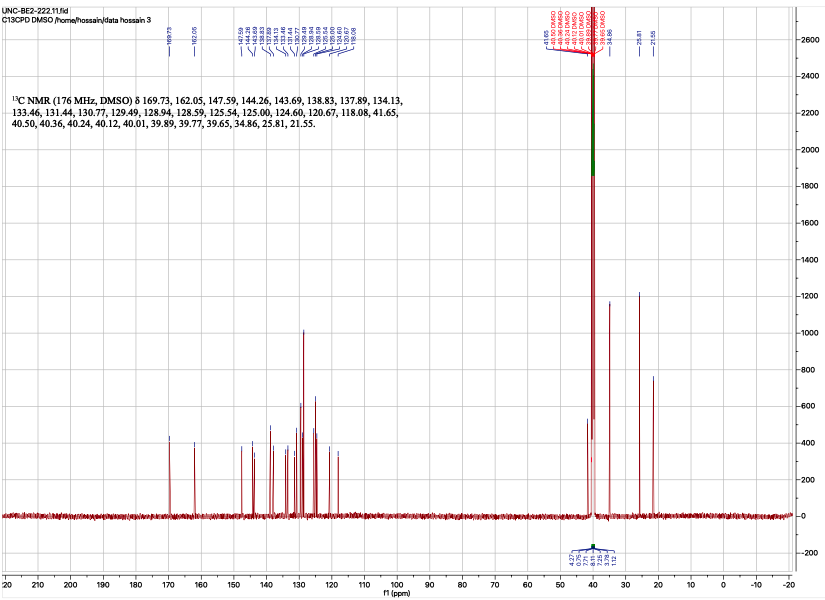

^1^H NMR for **6**

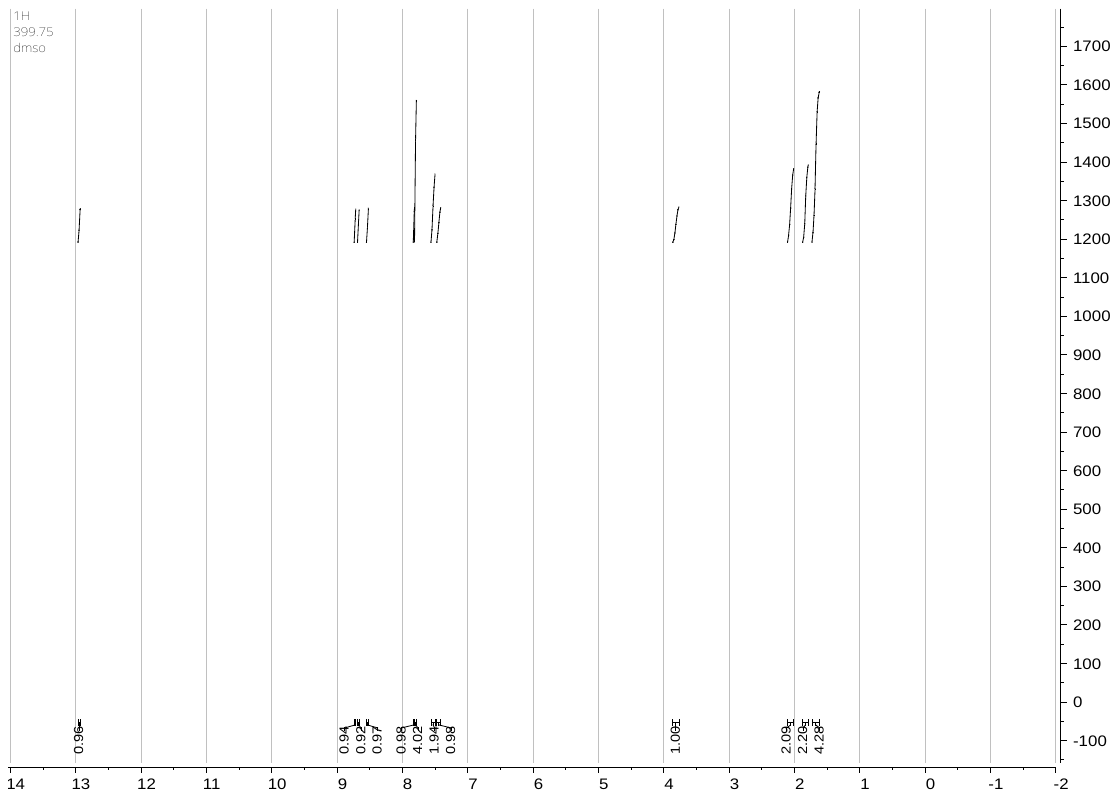

^13^C NMR for **6**

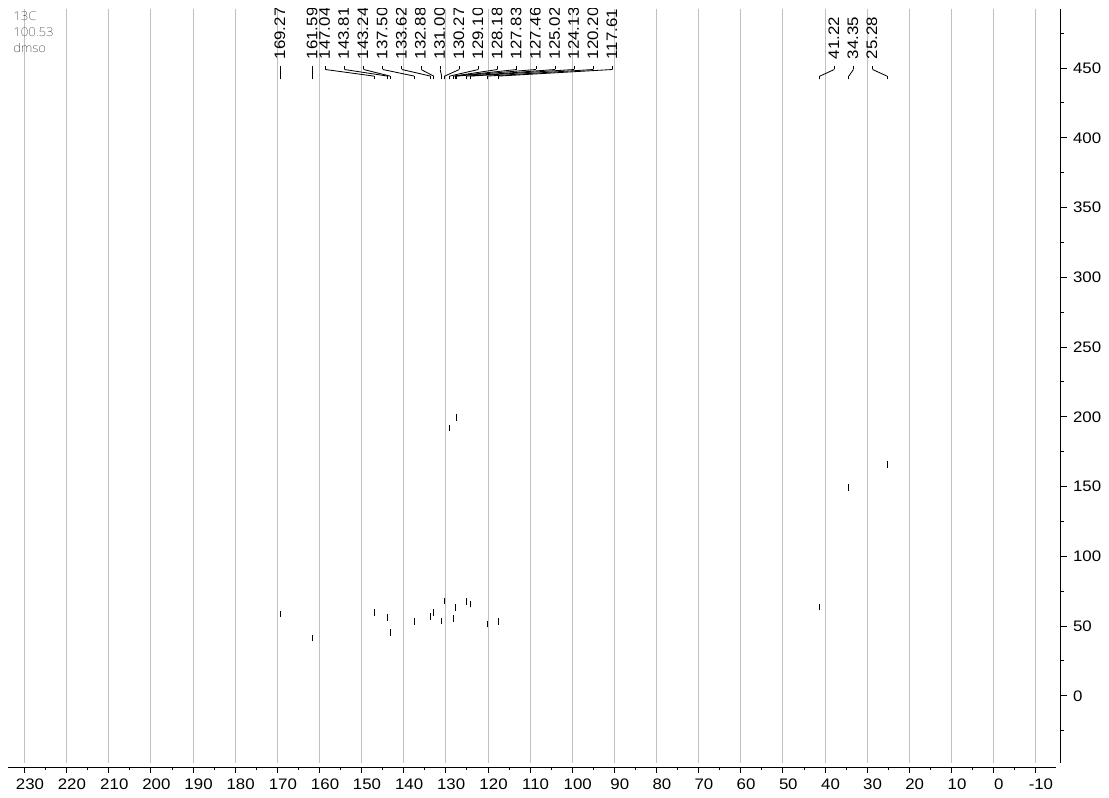

^1^H NMR for **7**

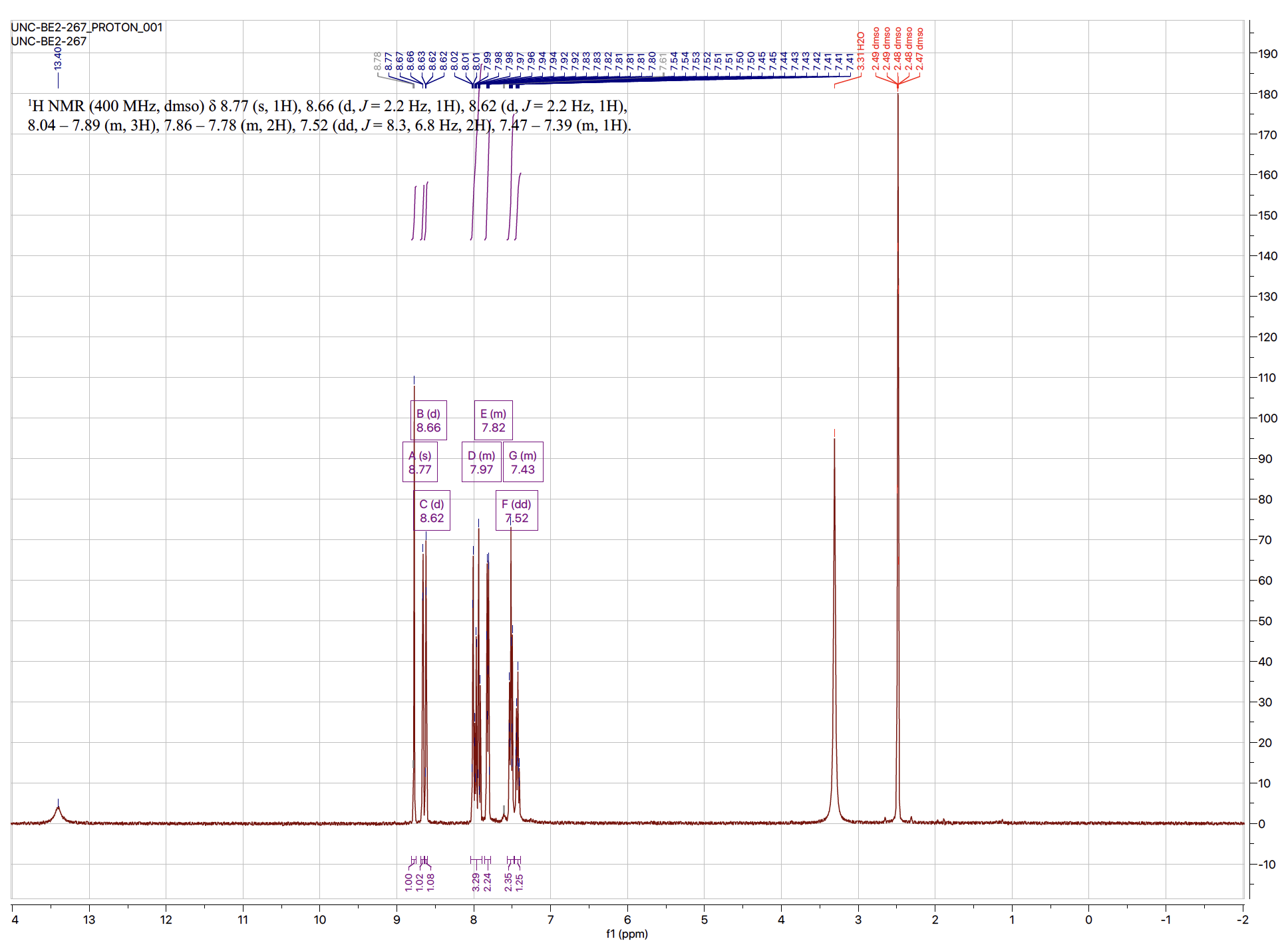

^13^C NMR for **7**

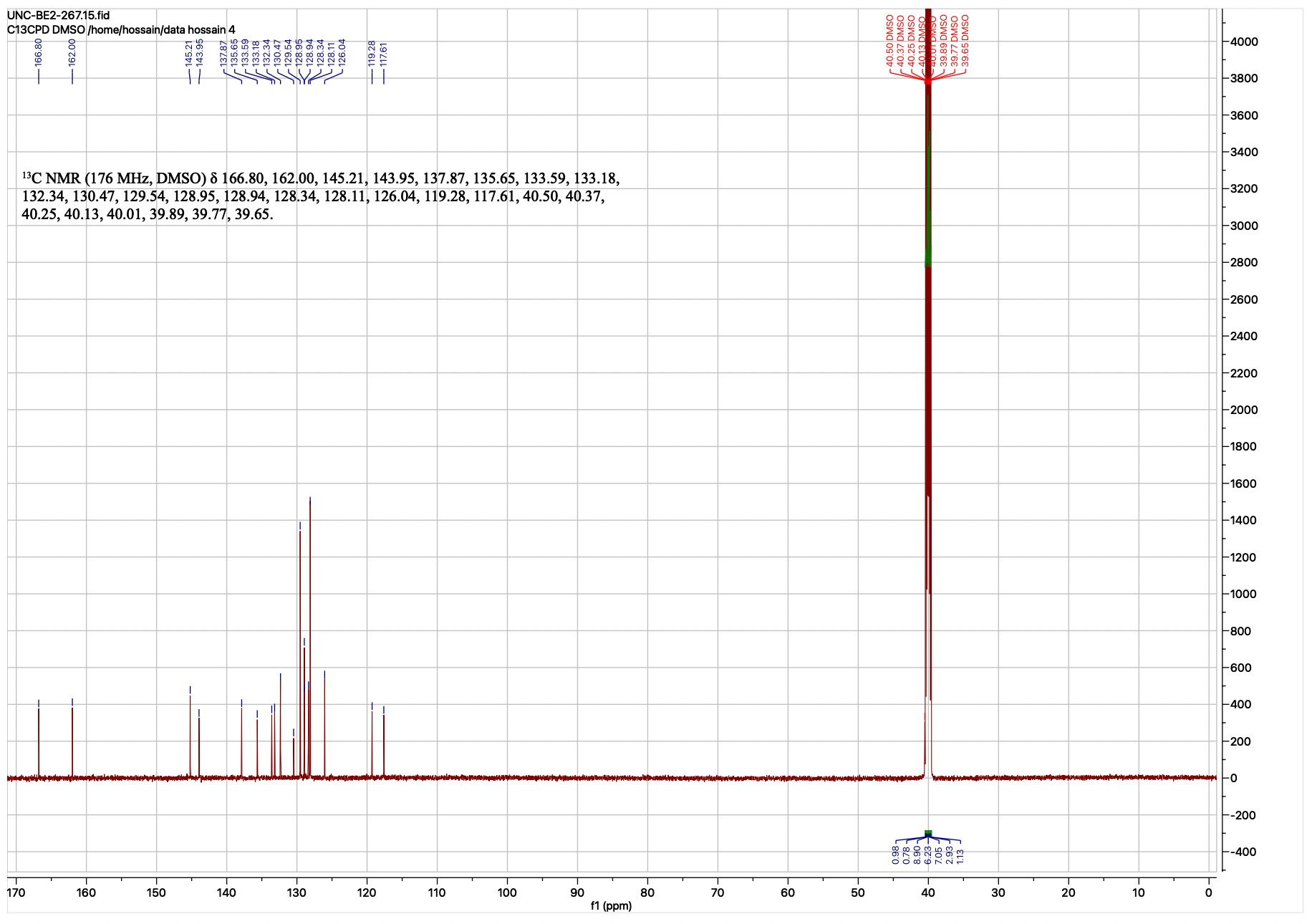

1.	Eduful, B.J.; O'Byrne, S.N.; Temme, L.; Asquith, C.R.M.; Liang, Y.; Picado, A.; Pilotte, J.R.; Hossain, M.A.; Wells, C.I.; Zuercher, W.J.; Catta-Preta, C.M.C.; Zonzini Ramos, P.; Santiago, A.S.; Counago, R.M.; Langendorf, C.G.; Nay, K.; Oakhill, J.S.; Pulliam, T.L.; Lin, C.; Awad, D.; Willson, T.M.; Frigo, D.E.; Scott, J.W.; Drewry, D.H. Hinge Binder Scaffold Hopping Identifies Potent Calcium/Calmodulin-Dependent Protein Kinase Kinase 2 (CAMKK2) Inhibitor Chemotypes. *Journal of medicinal chemistry* **2021**, *64*, 10849-10877, doi:10.1021/acs.jmedchem.0c02274.
